# TUMOR HETEROGENEITY AND STEMNESS CAN BE SHAPED BY MECHANO-STRUCTURAL PARAMETERS IN OSTEOSARCOMA

**DOI:** 10.64898/2026.05.22.727299

**Authors:** Zunaira Shoaib, Aleczandria S. Tiffany, Kyle Timmer, Brendan A.C. Harley, Joseph Irudayaraj, Timothy M. Fan

## Abstract

Chemoresistance and recurrence of osteosarcoma (OS) can be attributed to a subpopulation of OS cells known as OS Cancer Stem-like Cells (OS-CSCs). We hypothesized that certain mechano-structurally distinct niches within the bone and tumor might be more conducive to OS-CSC formation. Using biomaterial-based 3D scaffolds of distinct structural and mechanical properties (anisotropic soft ∼10 kPa vs isotropic stiff ∼35 kPa), we discovered that OS cells growing in softer, anisotropic scaffolds were rounder, softer, had higher stemness gene expression, showed increased chemoresistance to doxorubicin and cisplatin, and were more tumorigenic *in vivo.* Mechanistically, biological reprogramming of OS cells on these scaffolds occurred through niche-driven transcriptional changes and differences in chemoresistance-associated epigenetic pathways. Our study showed that a softer, anisotropic niche is more conducive for maintaining OS-CSCs and these findings could be translated to designing therapeutic strategies targeting heterogeneous tumor populations or modifying the microarchitecture to reduce CSC-favoring niches.

## INTRODUCTION

Primary bone tumor osteosarcoma (OS) is known to be chemoresistant with a high probability of recurrence at the site of resected primary tumor, with relapses occurring in 90% of surgery-only patients. Although adjuvant chemotherapy does improve survival of non-metastatic OS to 68%, only 15% patients survived with complete surgical remission. Moreover, 5-year survival decreases with recurrence, with only 16% surviving after the 2^nd^ recurrence (Gazouli et al., 2021). Unfortunately, there has been no significant improvement in the 5-year survival rate of metastatic OS over the past three decades, indicating a lack of understanding of the mechanism of OS metastasis and recurrence.

Although OS occurrence has no defined genetic driving factor, it is linked to bones undergoing remodeling with dynamically changing mechanical and structural properties which is evident by OS occurrence at age peaks related to rapid bone remodeling accompanied by structural and mechanical changes in bone. Additionally, patients of Paget’s disease are predisposed to OS, which is also characterized by similar bone remodeling and less dense and more brittle bone. Paget’s disease causes bone structural deformity due to high bone resorption characterized by increased number and size of osteoclasts, and decreased bone stiffness due to 14% decrease in Young’s modulus and 19% decrease in hardness of bone (Singer, 2016; Singer & Roodman, 2020). Moreover, Paget’s disease also worsens the prognosis of sarcomas as compared to sarcomas arising *de novo*, with the 5-year survival rate being 7.5% and 37% respectively (Ruggieri et al., 2010; Singer & Krane, 1998). Additional evidence of significance of biomechanics in OS is that the occurrence and prognosis of OS is bone location dependent, with greatest occurrence (42%) in the metaphysis of weight-bearing long bone, the femur, followed by the tibia (19 %) and humerus (10 %) (Jerome et al., 2009). In comparison, the site with the best prognostic outcome is the distal lower extremity in humans (Boerman et al., 2012). Earlier detection of tumors at certain sites might be one of the factors influencing this prognosis but it doesn’t fully explain why certain bone niches are more prone to occurrence and worse prognosis.

OS is proposed to arise from mutant MSCs or progenitor cells which are housed in the red bone marrow of the trabecular niche at the metaphysis. It is important to note that the metaphysis is made of anisotropic, trabecular bone, which is softer than cortical bone (Morgan et al., 2018). Both bone marrow and trabecular bone are highly heterogeneous in stiffness, with the range of marrow stiffness being 0.25 to 24 kPa (Jansen et al., 2015). Additional insight from bone metastasis of other cancers, such as pancreatic cancer, shows that tumor lesions in the bone are bone-region dependent, which leads to co-existence of both osteoblastic and osteolytic lesions derived from the same cancer cells in different regions of the same bone (Hirata et al., 2016). This is not surprising, as tumors have been reported to be heterogeneous at cellular, mechanical, and biochemical levels, with certain areas of the tumor being hypoxic or fibrotic, creating distinct biochemical and physical niches which might hinder access to therapies and anti-tumor immune cells (X. Li et al., 2019).

The cancer stem cell theory suggests that there might be a population of tumor-originating “cancer stem cells” (CSC) that lead to micro-metastasis in distant organs while remaining quiescent and chemoresistant, causing recurrence after the primary tumor has been removed (Oskarsson et al., 2014). Considering how metastatic tropism is seen in soft organs and how soft cells survive better in soft microenvironments; we can appreciate the importance of ‘niche’ mechanics in the cascade of cancer metastasis. The switch from a stem-like cell to a rapidly-dividing cell might be accompanied by a series of mechanobiological changes in the cell itself with the CSC having to attach or “anchor” to the scaffolding provided by fibrotic or stiff ECM, created through increased collagen crosslinking, consequently leading to clustering of integrins and Focal Adhesions (FAs), change in cytoskeletal structure, triggering cytoskeletal-linked mechanotransduction pathways, and increasing tumor lesion size (Levental et al., 2009; Shoaib et al., 2022). It is reported that cell attachment is essential for cell division, and differentiated cells are in turn stiffer than their progenitors (Darling et al., 2008). For the CSCs to survive during this entire process, they would have to be maintained in a CSC-conducive tumor microenvironment (TME) which might be different from the non-CSC TME due to the link between substrate mechanics (“suitable soil”) and cellular mechanics.

It has been established that mechanically different substrates inform cell behavior and plasticity (Discher et al., 2005). Whereas collagen alignment and anisotropy of tissue have been implicated in carcinoma metastasis (Conceição et al., 2024; Leng et al., 2021) and has shown to promote MSC growth in bone mimicking biomaterial scaffolds (Dewey et al., 2020). The anisotropy of trabecular bone in metaphysis, which is predisposed to OS occurrence, draws attention to its significance as an additional microarchitectural component to the mechanical component of the TME. The role of both structural (anisotropy vs isotropy) and mechanical (softness vs stiffness) properties of the bone TME in the plasticity, chemoresistance and tumorigenicity of OS is not well understood, and this study provides valuable insight into this topic by using mechano-structurally tunable, mineralized collagen scaffolds. Developed to promote regenerative healing of bone defects (Dewey et al., 2020; Ren et al., 2016), these scaffolds provide structural, compositional, and mechanical signals that can also be used to examine processes of OS activity. We hypothesize that certain niches within the bone might be more conducive for the maintenance and formation of OS CSCs than the niches harboring rapidly growing OS subpopulations. To test this hypothesis, we initially compared activity in two collagen scaffold variants, one with aligned pores and lower stiffness (A soft) and one with non-aligned pores and higher stiffness (NA stiff) to represent distinctly mechanically and structurally heterogeneous niches. We later expanded our studies to four variants (A soft, A stiff, NA soft, NA stiff) to explore the intermediate states between these two disparate niches.

## RESULTS

### Tunable 3D collagen scaffolds represented distinct mechano-structural bone niches

Two distinct 3D collagen scaffold variants were prepared by changing pore alignment, creating one with aligned pores and lower stiffness (A soft) and one with non-aligned pores and higher stiffness (NA stiff) (Figure 1 (A)). Distinct and reproducible mechanical properties of these mineralized scaffold variants were confirmed through compression tests (Figure 1 (B)). The two extreme conditions of A soft and NA stiff were selected for more detailed downstream analysis because they embodied more drastic and significant biological and mechano-structural differences. Compression tests showed a significant difference in elastic modulus of A soft (aligned pores/anisotropic and soft) and NA stiff (non-aligned pores/isotropic and stiff) scaffolds in all 3 conditions; when scaffolds were dry, when they were hydrated but unseeded (∼12.3 kPa and ∼31.2 kPa) and after being seeded with 143B cells (∼8.8 kPa and ∼26.7 kPa), with the A soft group being softer than the NA stiff (Supplementary Figure 1 (A)). Although hydration appeared to drastically soften the scaffolds but the difference between A and NA scaffolds remained significant. ESEM imaging and AFM of scaffolds showed that A and NA scaffolds were topographically different as well, as shown in Figure 1 (C-E).

**Figure 1.**
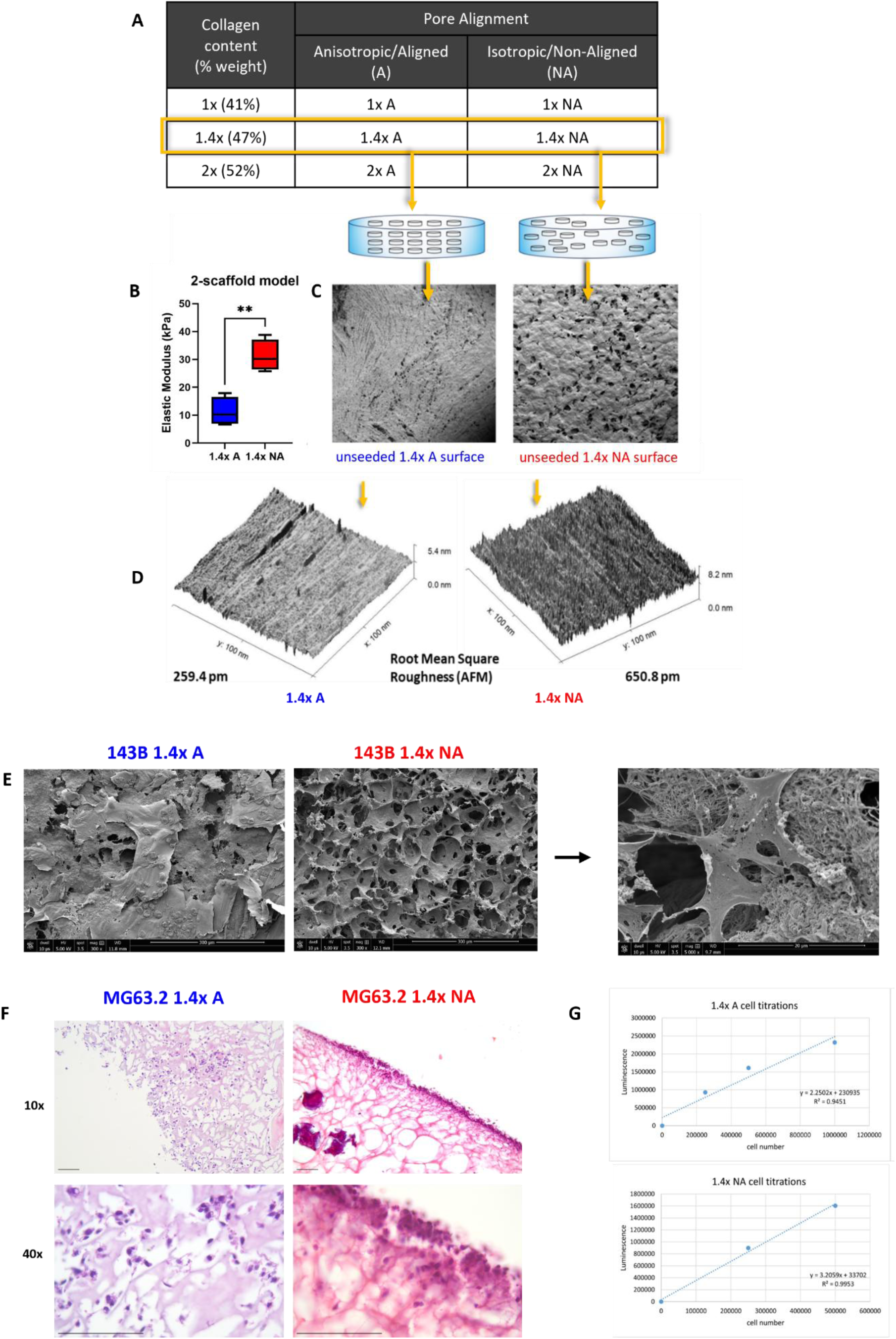
Production and characterization of 3D biomimetic scaffolds before and after osteosarcoma cell seeding. **(A)** The 2-scaffold model representing two distinct niches consisted of same collagen content (47%) and calcium-phosphate mineralization with variation in pore alignment from A (Aligned/Anistropic) (left) to NA (Non-Aligned/Isotropic) (right) **(B)** Measurement of stiffness (Young modulus) of the 2-scaffold models using compression tests (n=4) showed that NA was significantly stiffer than A. **(C)** SEM images of the top surface of unseeded, hydrated scaffolds. **(D)** AFM of top surface of dry scaffolds showed NA having significantly higher root mean square roughness as compared to A, corresponding to the modulus measurements. **(E)** SEM of scaffold surface seeded and populated with osteosarcoma (OS) cell lines; images at 300x magnification (scale bar=300um) and zoomed in image at 5000x magnification (scale bar=20um). **(F)** H&E staining of FFPE scaffolds populated with OS MG63.2 cells at 10x and 40x magnification showed difference in cell attachment and growth pattern between the 2-scaffold models (scale bar=200um). **(G)** Cell viability assay demonstrated biocompatibility and validity of our 2-scaffold models, with viability increasing in a cell concentration dependent manner.

### 3D collagen scaffolds were biocompatible and allowed OS growth

As the mineralized scaffolds are opaque, attached cells cannot be visualized by light microscopy after seeding. To visualize and verify if osteosarcoma cells had successfully attached and were growing in fabricated scaffolds, ESEM imaging, H&E imaging and viability assays of seeded scaffolds were performed (Figure 1 (E-G)). H&E imaging showed cells attached to the surfaces of and penetrated the seeded scaffolds in different patterns (Figure 1 (F), Supplementary Figure 1 (B)). alamarBlue and CellTiter-Glo assays showed that cells were metabolically active and that the luminescence signal from CellTiter-Glo increased in a cell number dependent manner (Figure 1 (G)).

### Softer, aligned niche (soft 1.4x A) upregulated stemness genes in OS and led to greater *in vitro* tumorigenicity

Higher expression of established stemness markers NANOG, OCT4, SOX2 were observed in aligned soft (1.4x A) vs non-aligned stiff (1.4x NA) 143B and MG63.2 cultures through real-time qPCR (Figure 2 (A)) after 4 days of culture. ALDH1A1, which has been reported to be a stemness gene in previous osteosarcoma studies (Mu et al., 2013), was also observed to be upregulated in 1.4x A cell population at both mRNA and protein levels (Figure 2 (A, B)). Previously, optimization experiments were carried out to demonstrate that day 4 of culture showed peak stemness gene expression (Supplementary Figure 1 (C)).

**Figure 2.**
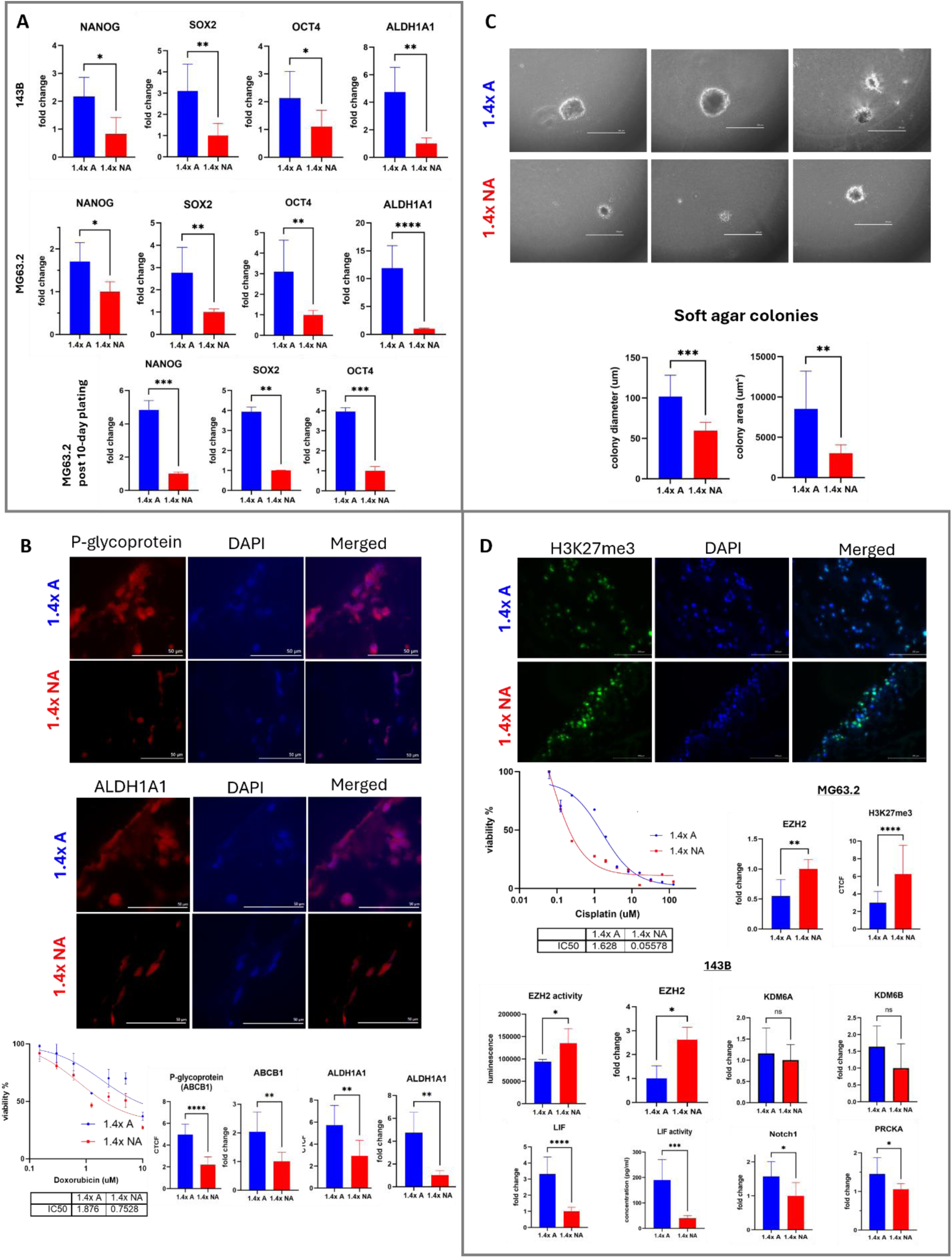
Cell populations in soft, aligned scaffolds (1.4x A) show cancer stem cell characteristics as compared to those in stiff, non-aligned scaffolds (1.4x NA). **(A)** Higher expression of hallmark stem cell genes NANOG, SOX2, OCT4, ALDH1A1 was observed in osteosarcoma cell lines (143B, MG63.2) after 4-day culture in 1.4x A soft, and this trend was conserved in cells harvested from scaffolds and grown on plates for 10 days. **(B)** OS 143B populations in 1.4x A soft were more resistant to Doxorubicin than 1.4x NA populations, which was linked to ABCB1/P-glycoprotein expression. 143B cells grown in 1.4x A soft niche showed higher protein expression of drug efflux pump P-glycoprotein and ALDH1A1 through immunofluorescence on FFPE scaffolds (scale bar=50um). Quantification of P-glycoprotein and ALDH1A1 immunofluorescence expression is shown in CTCF values (Corrected Total Cell Fluorescence). qPCR of ABCB1 and ALDH1A1 genes validate the higher expression of these factors in 1.4x A soft niche at both mRNA and protein levels. **(C)** 143B cells harvested from A soft cultures (represented as 1.4x A) grew bigger colonies in soft agar colony assay as compared to cells from NA stiff cultures (represented as 1.4x NA) (magnification 20x, scale bar=200um). Quantification and comparison of soft agar colony diameter and area (n=8-10) **(D)** Immunofluorescence of OS MG63.2 scaffold cultures show higher expression of epigenetic mark H3K27me3 in 1.4x NA stiff niche, which is linked to increased Cisplatin resistance (scale bar=100um). Higher expression of EZH2 mRNA and EZH2 activity was seen in MG63.2 and 143B 1.4x NA stiff cultures which correspond with the CTCF quantification of H3K27me3 showing significantly increased methylation in 1.4x NA stiff. Demethylase KDM6A/B expression did not show any significant difference between 2-scaffold models, indicating that methylation difference was due mainly to differential EZH2 expression and activity. Higher expression of downstream signaling pathways (PRKCA, Notch1, LIF) in EZH2-low 1.4x A soft scaffold cultures corresponded with increased Cisplatin resistance in 1.4x A soft niche.

Live cells harvested from 2-scaffold models were analyzed for their *in vitro* tumorigenicity by testing their spheroidal colony forming ability in soft agar. Populations harvested from soft 1.4x A resulted in bigger spheroidal colonies in soft agar than populations harvested from stiff 1.4x NA (Figure 2 (C)).

### OS population in softer, aligned niche (1.4x A soft) was more chemoresistant

Doxorubicin and cisplatin are two of the drugs used in the MAP regimen to manage osteosarcoma. More stem-like cancer cells (CSCs) are known to show higher chemoresistance. High chemoresistance against doxorubicin and cisplatin was observed in 1.4x A culture as compared to 1.4x NA. Mechanistically, resistance to these chemotherapies is acquired by OS cells through different pathways. Doxorubicin resistance has been attributed to an upregulation of drug efflux pump ABCB1 (p-glycoprotein) (Mirzaei et al., 2022). In alignment with this finding, 1.4x A cell population showed an increase in ABCB1 transcription and expression of the encoded P-glycoprotein, along with higher doxorubicin resistance (Figure 2 (B)).

### Niche mechanics influenced chemoresistance-linked methylation

Cisplatin resistance in the OS CSC population has been linked to low expression of methyltransferase EZH2 and its methylation marker H3K27me3. This change in methylation status triggers multiple pathways as demonstrated in the study by He et al. (He et al., 2019), which have been summarized in the schematic (Supplementary Figure 1 (D)).

Accordingly, to further verify change in methylation as a stemness parameter, a preliminary experiment was carried out using parental non-metastatic and derived metastatic OS cell lines (HOS, MNNG-HOS, 143B), which showed almost negligible transcriptional and proteomic expression of EZH2 and H3K27me3 in the most malignant and highly metastatic cell line 143B as compared to less aggressive MNNG-HOS and non-metastatic HOS (Supplementary Figure 1 (D)).

The cell population in 1.4x A showed a decrease in EZH2 transcription and EZH2 methylation activity as compared to 1.4x NA cultures (Figure 2 (D)). In accordance with the findings of He et al. (He et al., 2019), there was an increase in LIF transcription and LIF release in the media from the more CSC-like population in 143B 1.4x A soft scaffolds (Figure 2 (D)). There was an increased expression of Notch1 and PRCKA in 1.4x A soft population as compared to 1.4x NA. Immunofluorescence of FFPE scaffolds showed decreased expression of H3K27me3 in 1.4x A soft populations as compared to 1.4x NA stiff populations (Figure 2 (D)), corresponding to decreased EZH2 activity in 1.4x A soft population.

### Niche properties drove differential gene expression in RNA-Sequencing

RNA-sequencing showed niche-driven differences in gene expression. Using the FDR p-value threshold of 0.3, a total of 2,845 genes were significant for at least one of the pairwise comparisons or the interaction term (Figure 3). Most importantly, there were 87 genes with the same magnitude and direction of response to scaffold type in both cell lines. Gene targets were narrowed down from the RNA Seq data (SNHG1, KLF6, KLF6-SV1) due to their high fold change across cell lines and implication in stemness pathways.

**Figure 3.**
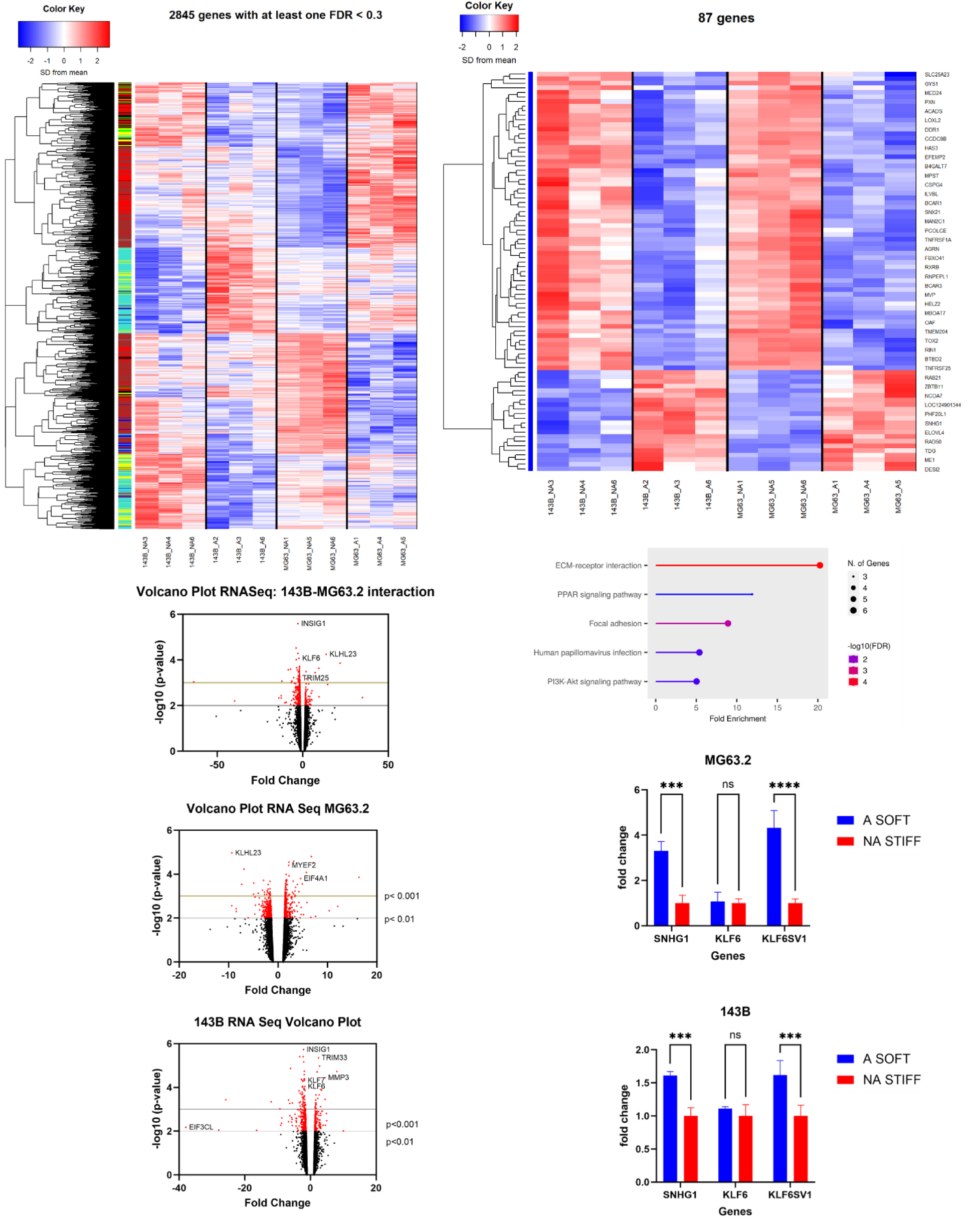
RNA-Sequencing showed niche-driven differences in gene expression across cell lines. Differential gene expression between A and NA cultures in 143B and MG63.2 shown per cell line (top left). 87 genes with common direction and magnitude of difference between A and NA cultures were seen across both cell lines (top right). Data showed enrichment in genes belonging to the PPAR pathway, PI3K-Akt pathway, and cell-substrate interactions. Volcano plots showing RNA Seq data represented as fold change of gene expression, with genes with most magnitude change per cell line highlighted (bottom left). qPCR of targets picked from RNA Seq data (SNHG1, KLF6, KLF6-SV1) confirmed the higher fold change of stemness related gene SNHG1 in soft 1.4x A cultures, whereas only the oncogenic KLF6-SV1 splice variant of KLF6 splice was upregulated in soft 1.4x A cultures, not the full KLF6 variant.

Although RNA Seq data showed high fold change in KLF6, it does not discriminate between its different isoforms and variants. It is important to note that one of the KLF6 isoforms called isoform D or variant KLF6-SV1, has been classified as an oncogene by recent studies (Hu et al., 2021). Targeted qPCR of KLF6 and KLF6-SV1 done independently for both cell lines 143B and MG63.2, showed that it was the KLF6-SV1 variant that was significantly enhanced in A soft (1.4x A) populations as compared to NA stiff (1.4x NA) populations (2-way ANOVA with Sidak’s multiple comparison, p<0.001), whereas KLF6 showed no significant differences (Figure 3). Furthermore, SNHG1 showed a similar trend of being increased in A soft (1.4x A) populations as compared to NA stiff (1.4x NA) populations (2-way ANOVA with Sidak’s multiple comparison, p<0.001) and complemented the RNA Seq data (Figure 3).

### Niche properties drove cytoskeletal organization and YAP nuclear localization

Integrins and YAP/TAZ play a significant role in sensing substrate stiffness and regulating mechanotransduction-driven cell plasticity respectively. First, transcriptional analysis revealed that levels of YAP, TAZ, and Focal Adhesion Kinase (FAK) were significantly increased in populations of 1.4x NA stiff niche as compared to 1.4x A soft niche (Figure 4 (A)). Contrastingly, Integrin Alpha 6 (ITGA6) gene was upregulated in 1.4x A soft scaffolds, while Integrin Beta 1 (ITGB1) showed no transcriptional difference across niches (Figure 4 (A)). Immunofluorescence of FFPE sections of 1.4x A soft and 1.4x NA stiff scaffolds showed a change in OS cell morphology, actin arrangement and active YAP nuclear levels (Figure 4 (B)). As expected, the stretched, elongated 143B OS cells in 1.4x NA stiff niche had higher active YAP levels in the nucleus which is indicative of a more differentiated phenotype. Whereas the cell population growing on softer, aligned matrix of 1.4x A scaffolds showed rounder cells with lower active YAP in the nucleus, representing a more stem-cell like phenotype.

**Figure 4.**
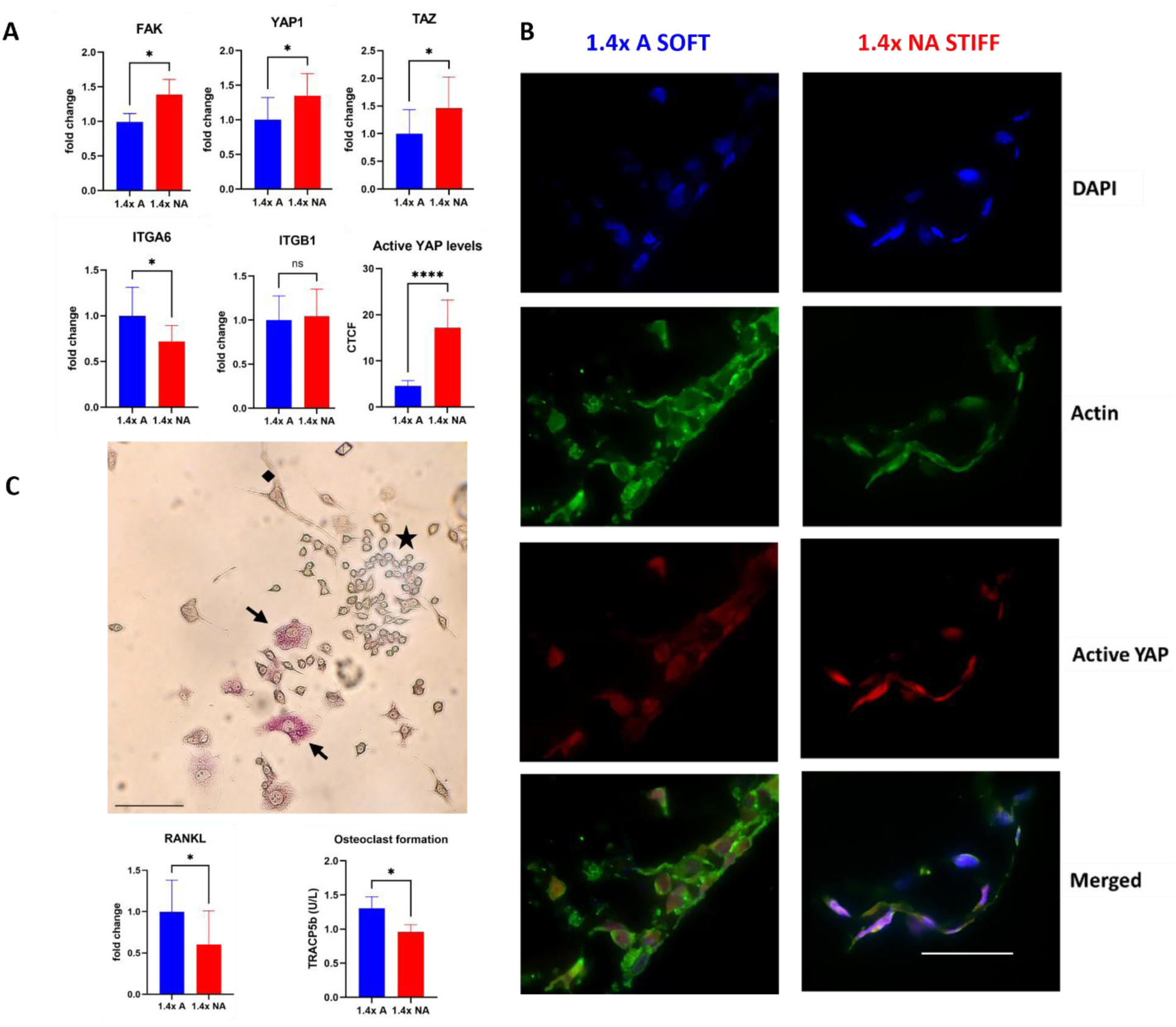
Mechano-structural parameters drive cytoskeletal arrangement, YAP nuclear localization and osteoclast formation. **(A)** Focal Adhesion Kinase (FAK), YAP and TAZ were increased at transcriptional level in 1.4x NA stiff cultures. Quantification of active YAP protein levels in the nucleus from FFPE immunofluorescence data shown in terms of Corrected Total Cell Fluorescence (CTCF), confirmed significantly higher YAP activity in 1.4x NA stiff niche as compared to 1.4x A soft niche. Integrin Alpha 6 gene (ITGA6) was upregulated in 1.4x A soft niche but no difference was seen in Integrin Beta 1 (ITGB1) gene expression. **(B)** Active YAP is important for mechanotransduction of signals sensed from the substrate surface to regulate osteogenic differentiation. Immunofluorescence of FFPE stained scaffold sections showed a distinct change in cytoskeletal arrangement and cell shape across the two niches, with a rounder phenotype and decreased nuclear YAP in 1.4x A soft niche cultures as compared to a stretched, elongated phenotype in 1.4x NA stiff niche with higher nuclear YAP levels, indicative of a more stem-like and less stem-like phenotype, respectively (magnification 60x, scale bar=50um). **(C)** Co-culture of OS 143B cells (black diamond) with macrophages/pre-osteoclasts (black star) induces the formation of macrophages into active osteoclasts (black arrows), which can be detected by TRACP 5b purple staining under a light microscope on day 4 of co-culture (scale bar=200um). Increased RANKL transcriptional levels in 1.4x A soft niche corresponded with increased osteoclast formation in 1.4x A soft niche as indicated by increased TRACP5b levels measured through ELISA.

### Softer, aligned niche (soft 1.4x A) showed greater osteoclast formation

RANKL released from OS cells bind to RANK receptors on pre-osteoclasts such as macrophages to induce their terminal differentiation into active osteoclasts, which are responsible for bone resorption during the bone remodeling process in OS. Active osteoclasts contain and secrete TRACP5b (Tartrate-resistant acid phosphatase 5b) which differentiates them from macrophages which only release TRACP5a. Preliminary experiment optimizing co-culture of OS and macrophages on petri plates showed successful active osteoclast formation on day 4 of co-culture, as indicated by purple staining of TRACP5b in mature osteoclasts (Figure 4 (C)). Then TRACP5b levels in scaffold culture media was used as a readout for osteoclast formation in scaffolds co-cultured with osteosarcoma and macrophage cells. Increased RANKL levels in co-cultures grown on 1.4x A soft niche corresponded with increased TRACP5b release in 1.4x A soft niche as compared to 1.4x NA stiff co-cultures, indicating that A soft niche was more conducive for higher osteoclast formation (Figure 4 (C)).

### Cell population in softer, aligned niche (A soft) was softer than in stiff, non-aligned niche (NA stiff)

AFM tests showed a significant difference in elastic modulus of 143B cells grown on soft 1.4xA and stiff 1.4xNA scaffolds. Both live (1 kPa) and fixed (6.2 kPa) 143B cells growing on softer, 1.4xA scaffolds were around 4 times softer than live (4 kPa) and fixed (24.3 kPa) 143B cells growing on stiffer, 1.4xNA scaffolds (Figure 5). Height and adhesion maps were used to confirm that the area being scanned is a cell (Supplementary Figure 2). Force map corresponds to height map, with areas of increased height representing the cell. Similarly, height and Zsensor retrace mark out the cell on surfaces, although as the scaffold surface is softer and more heterogeneous than plastic surface, the cell outline is not as sharp.

**Figure 5.**
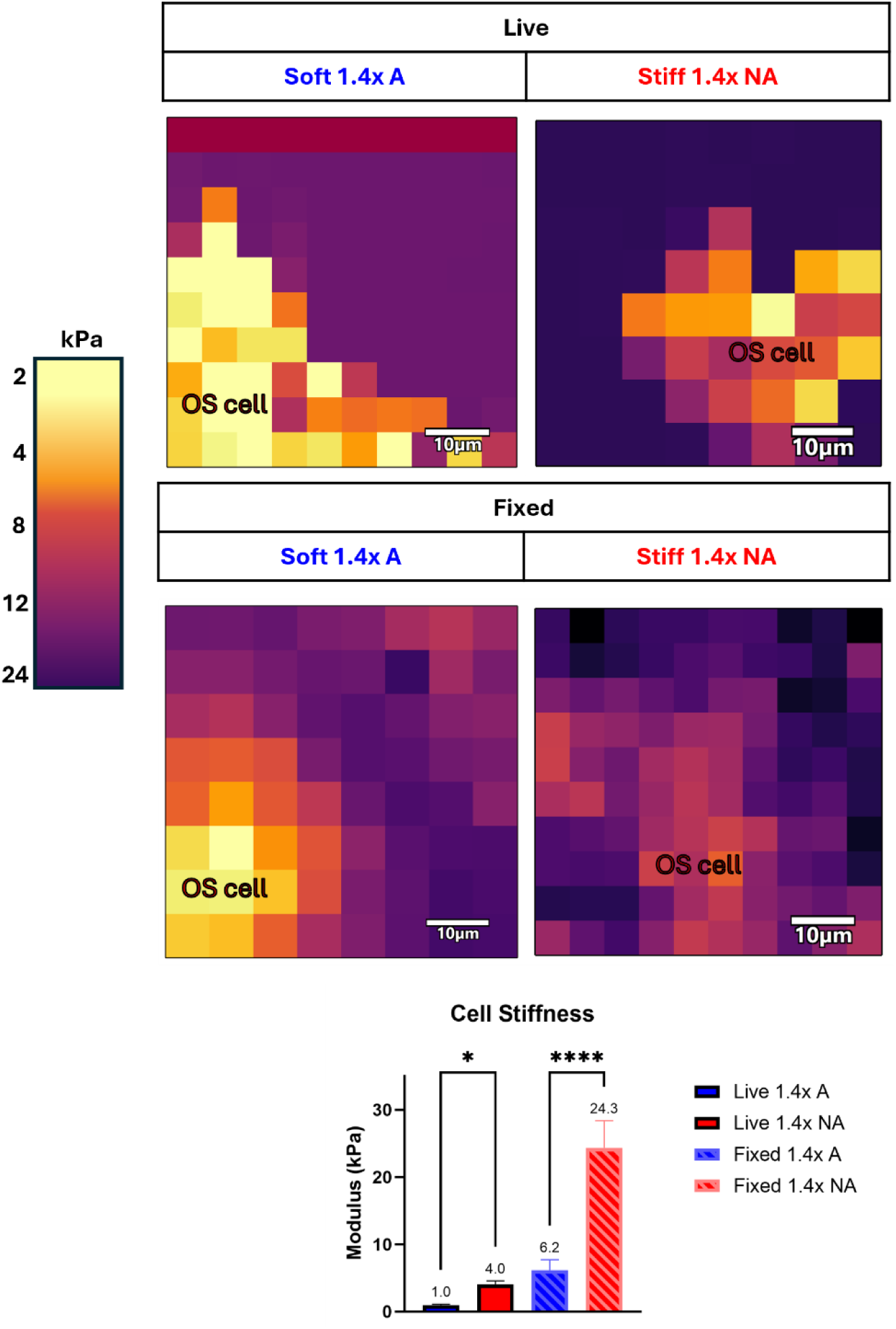
AFM analysis show OS cells grown on soft 1.4x A scaffolds are softer than cells on stiff 1.4x NA scaffolds, in both live and fixed conditions. Scale bar and image coloring fixed to highlight the cell on scaffold surface and to show comparative differences in cellular stiffness, calculated and shown as Young’s Modulus (kPa) in the graph.

### Niche mechano-structural properties in expanded 4-scaffold models drove stemness gene expression and *in vitro* tumorigenicity

We expanded our studies from two variants (A soft, NA stiff) to four variants (A soft, A stiff, NA soft, NA stiff), also referred to as the 4-scaffold model, to explore the intermediate states between these two distinct niches and to evaluate any matrix effect between stiffness and alignment. Each group in the 4-scaffold model was structurally (anisotropic ‘A’ v isotropic ‘NA’) and mechanically (soft vs stiff) distinct as confirmed through compression tests (Figure 6 (A)).

**Figure 6.**
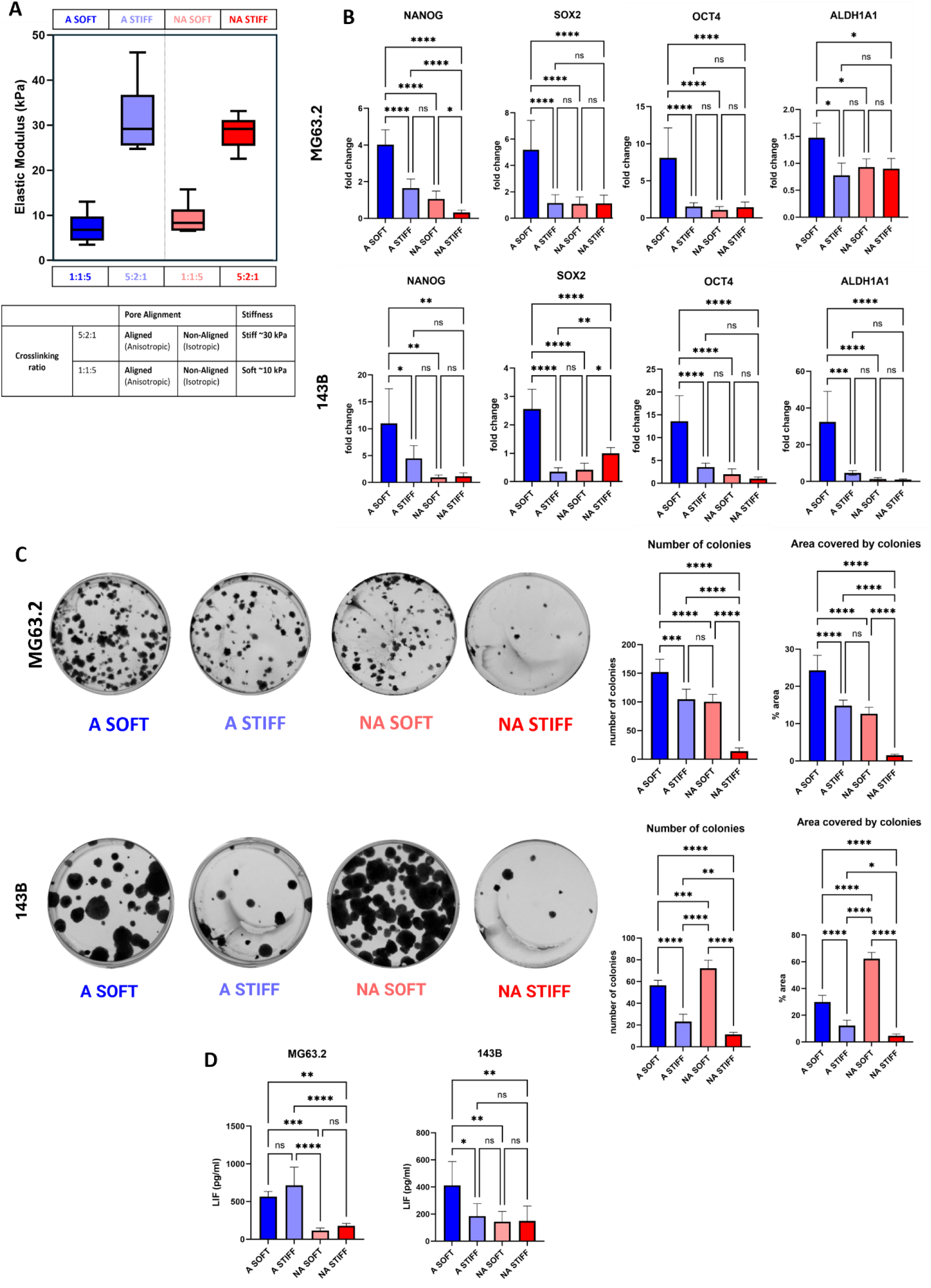
Osteosarcoma cells show increased cancer stem cell properties on ‘A soft’ scaffolds among the 4-scaffold models. The 4-scaffold models represent the range between 1.4xA (A soft) and 1.4xNA (NA stiff) 2-scaffold models, adding the intermediate forms of A stiff and NA soft to the pre-existing niche models. **(A)** compression tests of hydrated, unseeded 4-scaffold models show the same Young Modulus of stiff (∼35 kPa) and soft (∼10 kPa) models, but with different alignments (A or NA). **(B)** Both 143B and MG63.2 cultures in the 4-scaffold models show highest stem cell gene expression in A soft niche, which is representative of 1.4x A soft in the 2-scaffold model (ANOVA and Tukey’s multiple comparisons across groups). **(C)** Colony assay show highest tumorigenic potential of cells harvested from soft models, with MG63.2 showing highest colony number and area in A soft populations, and 143B showing highest colony number and area in NA soft, followed by A soft. Populations from stiff scaffolds show lowest colony formation across both cell lines, but A stiff shows higher colony formation and tumorigenic potential than NA stiff, showing an influence of alignment in stiffer niches. **(D)** Highest release of IL6-like cytokine LIF was observed in A soft niche in 143B 4-scaffold models, whereas in MG63.2 seeded models, both anisotropic scaffolds (A soft, A stiff) showed significant increase over isotropic (NA soft, NA stiff) scaffolds, indicating that the role of anisotropy and stiffness might be more complex than originally thought.

qPCR analysis of stemness genes carried out after following the same protocol for cell culture and RNA extraction as the 2-scaffold model showed similar trends as seen in the 2-scaffold model. A soft, corresponding to 1.4x A, still showed the highest expression of stemness genes, whereas NA stiff, corresponding to 1.4x NA, showed the lowest expression of stemness genes across the 4 groups (Figure 6 (B)).

Live cells harvested from 4-scaffold model cultures were analyzed for their colony forming ability by growing them in standard tissue-culture treated plastic 6-well plates. Colony assays from scaffold-harvested MG63.2 cells showed the highest colony numbers and area covered in the A soft group, whereas scaffold-harvested 143B cells showed the highest colony numbers and area covered in the NA soft group (Figure 6 (C)). Overall, soft scaffold-derived populations showed greater colony formation than stiff scaffold-derived populations.

Increased transcriptional and secreted levels of LIF seen from the more CSC-like population in 143B 1.4x A soft scaffold in the 2-scaffold models (Figure 2 (D)), were also consistent in the 4-scaffold models (Figure 6 (D)). However, in MG63.2 seeded 4-scaffold models, alignment played an unexpected role, and we observed high levels of LIF released from both A soft and A stiff niche cultures as compared to NA soft and NA stiff (Figure 6 (D)). This indicated there might be matrix effect created by anisotropy and stiffness, and their role in LIF pathway might be more complex.

### Soft scaffold-derived cells showed greater *in vivo* tumorigenicity

MG63.2 and 143B populations from 4-scaffold models (A soft, A stiff, NA soft, NA stiff) and 2-scaffold models (A soft, NA stiff) were harvested after 4 days of growth and mixed with Matrigel to create para-tibial tumors in nude mice (n=4). After 12 weeks, one of the mice from the NA soft group had to be euthanized because it had reached 2 mm tumor diameter. At the defined time point, all mice were euthanized and primary tumor growth at the right tibia and pulmonary metastases were analyzed. Following H&E staining, pathological analysis and review was kindly done by pathologist Dr. Keith Bailey. Interestingly, in MG63.2 mice, NA soft group showed the highest growth, although equal penetrance was seen in NA soft and A soft (Figure 7 (A)). A stiff group showed higher penetrance than NA stiff group, showing that anisotropy might have an influence on stiffer population. Pulmonary micro-metastases were seen only in NA soft.

**Figure 7.**
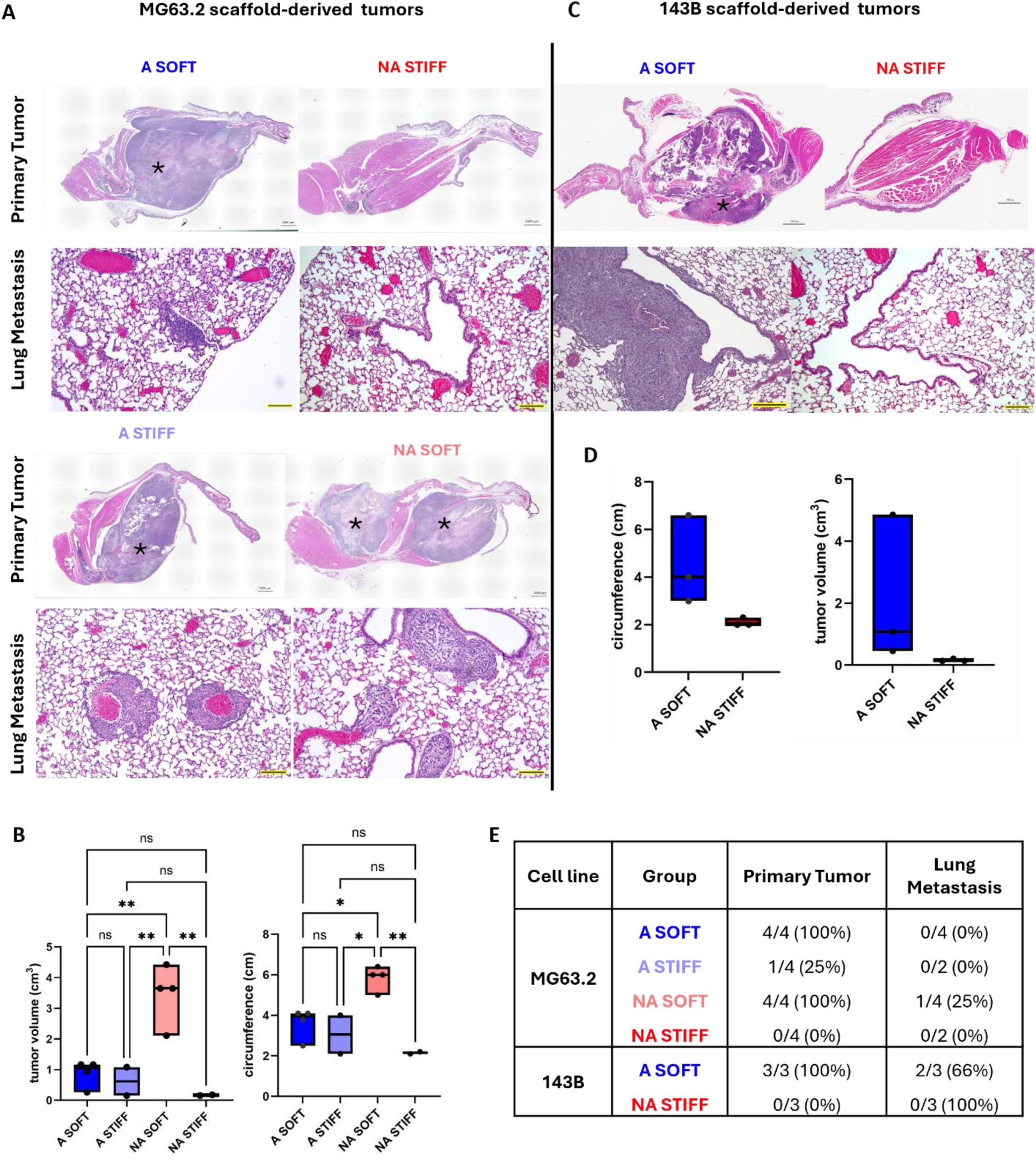
Soft scaffold-derived osteosarcoma cells show highest tumorigenicity *in vivo*. **(A)** Images of right tibia and corresponding histology with H&E stain shows para-tibial tumors derived from scaffold-harvested MG63.2 cells, with visible primary tumors (black star) in all groups except NA stiff (scale bar=2000um). Corresponding lung histology from each group is shown (magnification 10x, scale bar=200 um). **(B)** Quantification of tumor circumference (cm) and volume (cm^2^) from MG63.2 scaffold-derived tumors shown in graphs. **(C)** Images of right tibia and corresponding histology with H&E stain shows para-tibial tumors derived from scaffold-harvested 143B cells, with visible primary tumor (black star), showing greater tumor penetrance in A soft over NA stiff, (scale bar=2000 um). Corresponding lung histology shows pulmonary micro-metastases in A soft group but not in NA stiff group (magnification 10x, scale bar=200 um). **(D)** Quantification of tumor circumference (cm) and volume (cm^2^) from 143B scaffold-derived tumors showed in graphs. **(E)** Percentage of tumor penetrance and pulmonary micro-metastases across scaffold-derived tumor groups have been summarized in the table.

In a separate experiment, 143B populations from the 2-scaffold model (A soft, NA stiff) were harvested after 4 days of growth and mixed with Matrigel to create para-tibial tumors in nude mice (n=3). At the 60-day time point, all animals were sacrificed, and the right tibia and lungs were harvested and processed for H&E staining (Figure 7 (B)). All mice in the A soft group showed successful xenograft tumors (100% penetrance), whereas there was no growth in NA stiff group. Histological analysis showed metastases in lungs of A soft group only (Figure 7 (B)).

Overall, soft populations showed higher tumorigenicity *in vivo* as summarized in Figure 7 (C), but how isotropy or anisotropy was influencing this *in vivo* behavior warrants further investigation.

A schematic summary of this study is shown in Figure 8, which provides an overview of the experimental design.

**Figure 8.**
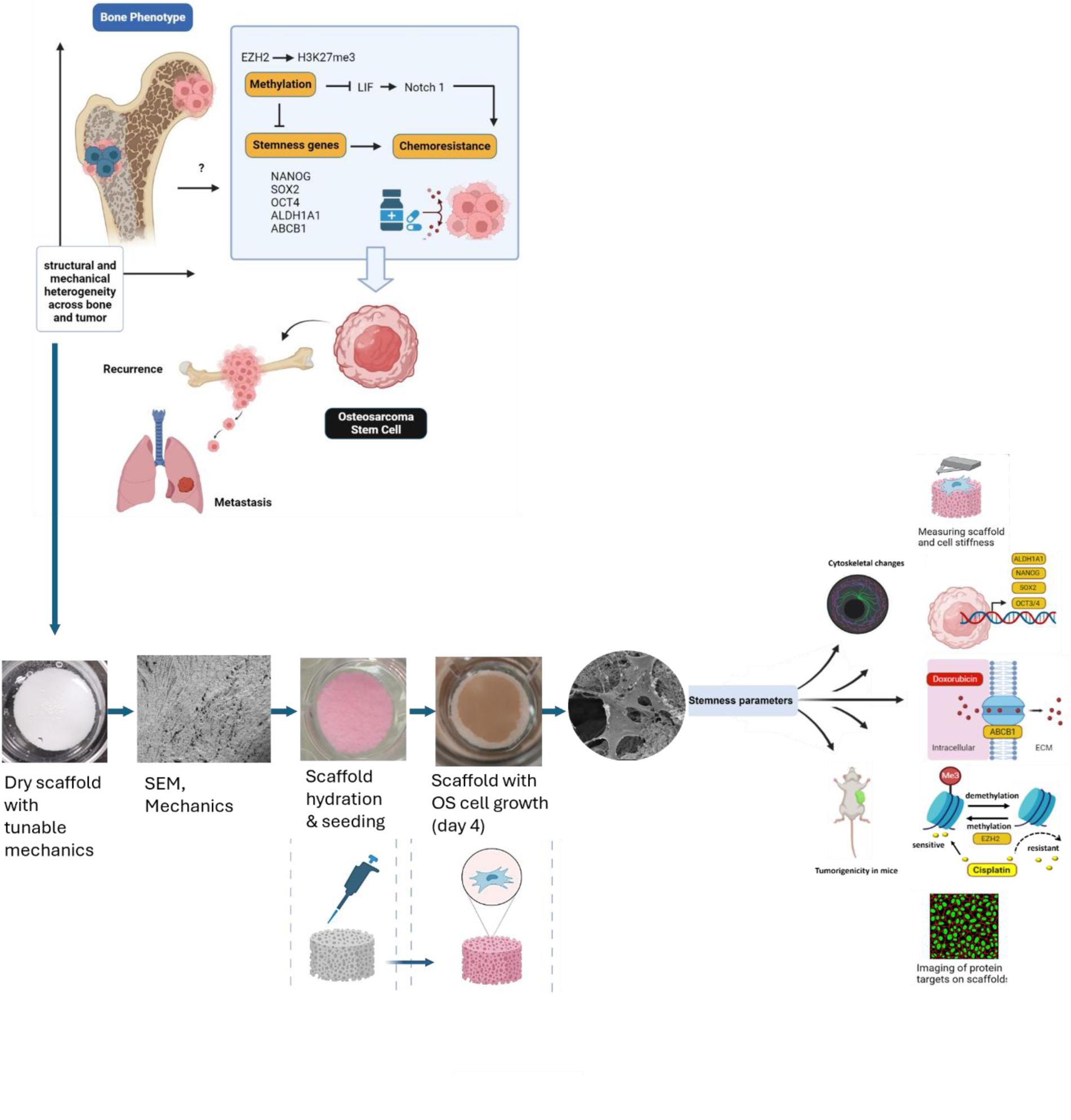
Schematic of study design.

## DISCUSSION

Considering pre-disposition of OS in bones undergoing structural and mechanical changes, as well as the mechanically distinct niches within the tumor, we hypothesized that structurally and mechanically distinct bone niches might be housing distinct tumor subpopulations, and certain niches might be more conducive for OS CSC formation and maintenance. We report that just changing the mechanical TME alone induces reprogramming in OS biology, enriching OS CSCs in the softer, anisotropic niche.

Studies investigating cellular mechanics have shown that substrate stiffness can induce lineage-specific differentiation or maintain stemness (Engler et al., 2006). There is contradictory evidence on how stiffness might influence tumor cells. Carcinomas and sarcomas might behave differently, and CSCs might behave differently from other tumor populations still, with breast CSCs preferring substrate stiffness of 5 kPa while OS cells showing greater survival on 50-55 kPa (Dailiana, 2008; Jabbari et al., 2015). Another study showed that OS cells grown on softer 1 kPa gels showed increased chemoresistance and higher stemness gene levels than cells on stiffer 20-55 kPa gels (S. Li et al., 2020). Most cancers show metastatic tropism to soft organs such as the lung, liver, brain, and bone marrow, which have a tissue stiffness in a range of 0.1 – 1 kPa (Valastyan & Weinberg, 2011), in addition to a growing body of evidence showing that softer substrates might be more conducive for the formation and maintenance of cancer stem cells (Deng et al., 2022).

Based on the literature, we created 3D collagen biomaterial models within the range of stiffness 10 – 40 kPa by changing the amount of crosslinking and tested stemness parameters of cell populations in each niche. We first validated that our 3D mineralized collagen models are conducive for OS cell attachment and growth by showing that cell viability on the scaffolds increased in a cell number dependent manner. ESEM and H&E imaging further confirmed that OS cells can successfully attach to, grow on and penetrate in the collagen scaffolds.

Stem-like cells are expected to express hallmark stemness genes like SOX2, NANOG, OCT4 and are shown to be softer and rounder than more differentiated, rapidly dividing cancer cells. Initially, hallmark stemness genes (NANOG, SOX2, OCT4) were evaluated at different time points to assess peak expression in scaffold cultures, which was concluded to be on day 4 of culturing. OS populations grown in soft scaffold cultures (∼10 kPa) with anisotropic/aligned pore structure (A soft) consistently showed an increased expression of hallmark stemness genes as compared to populations grown in stiffer scaffolds (∼30 kPa) with isotropic/non-aligned pore structure (NA stiff). This trend remained consistent even after expanding the 2-model system to a 4-scaffold model. Cells growing on A soft were 4 times softer than the ones on NA stiff, in both live and fixed conditions as determined by fluid force-mapping by AFM. This corresponds with the overall idea that cell stiffness is also dependent on substrate stiffness as human OS cells were shown to be significantly softer on collagen-coated surfaces as compared to glass substrate in another study (Docheva et al., 2008). It also aligns with previous studies showing that stem and progenitor cells are softer than differentiated cells (Darling et al., 2008) and tumorigenicity of cells is linked to its mechanical properties, as cancer cells are known to be softer than normal cells and CSCs are softer than other cancer subpopulations (Chen et al., 2022; Lin et al., 2015). This makes sense considering that metastasizing cells would need to be softer and flexible when extravasating across barriers and traveling via the narrow vessels of the circulatory system, a phenomenon coined as “mechanoadaption” (Rianna et al., 2020). The study by Rianna et al (Rianna et al., 2020) reported that osteosarcoma U20S cells underwent cytoskeletal changes and a decrease in stiffness from 5.6 kPa to 2.1 kPa as they moved through confined 3D matrices. Moreover, this study aligned with our findings that YAP nuclear localization triggered in stiffer cells growing on stiffer substrates and that softer cells do not show YAP nuclear localization in conjunction with cytoskeletal re-arrangement.

Some studies dissecting OS mechanobiology have concluded that a stiffer TME is more conducive for OS growth than softer TME and this has been equated to OS stemness. However, these studies showed that mechanical force and stiffness caused increased OS proliferation due to increased FA formation and integrin clustering, leading to enhanced ECM-integrin downstream signaling (Adamopoulos et al., 2017) which makes sense considering the influence of stiffness on cell growth. Based on a previous report that a stiff ECM induces YAP nuclear translocation and activation, leading to transcription of differentiation-related genes and differentiation of MSCs into osteoblasts (Dupont et al., 2011), we evaluated the expression of YAP/TAZ downstream of cytoskeletal changes. In alignment with the previous reports (Dupont et al., 2011; Rianna et al., 2020), we observed significant changes in actin organization and cell shape accompanied with increased YAP transcript and YAP nuclear localization in the stretched, elongated cells growing on stiff 1.4xNA, as compared to rounder, softer cells on soft 1.4xA, further confirming the expected phenotype of a more differentiated versus a more stem-like populations on these niches respectively.

Moreover, YAP/TAZ activity was reportedly required for osteogenic differentiation from progenitor cells Tang et al., 2016), highlighting the link between mechanotransduction and cell plasticity. Multiple studies have associated increased YAP/TAZ nuclear translocation in OS with differentiation and increased cell proliferation (Dupont et al., 2011; Luu & Viloria-Petit, 2020), which has made it a marker for OS aggression. However, these studies don’t account for heterogenous tumor subpopulations. One study has linked this behavior to stemness (Basu-Roy et al., 2015) by showing that SOX2 downregulates the Hippo pathway and upregulates YAP. However, another study found no link between ITGB/YAP/TAZ and chemoresistance, while still maintaining YAP/TAZ as a prognostic factor in OS (Bouvier et al., 2016). In our study, Integrin B1 (ITGB) expression did not show any changes between the two soft and NA stiff populations. Our findings suggest that YAP/TAZ could be a marker for a more differentiated, less quiescent subpopulation found in a stiff ECM, and in that context, it would make sense to consider YAP/TAZ as a marker for OS aggression and increasing tumor burden, instead of a marker for CSCs. Interestingly, we saw YAP decrease at both transcriptional and protein levels along with increased RANKL and increased active osteoclast formation in A soft co-culture, which agrees with previous reports that showed that RANKL phosphorylates and downregulates YAP levels during osteoclastogenesis (Takito & Nakamura, 2020).

We used histone methylation as another parameter for stemness based on existing evidence that links substrate stiffness to histone methylation and cancer cell stemness. A study by Tan et al. (Tan et al., 2014) showed that a softer substrate of 90 Pa decreased histone methylation at H3K9, while a stiffer matrix of 1 kPa showed increased methylation. This stiffness-dependent decreased methylation led to increased stemness as measured by enhanced SOX2 expression in melanoma cells. Consistent with the seminal studies of He et al. (He et al., 2019) and Lu et al. (Lu et al., 2020), populations from softer, anisotropic scaffolds (1.4x A soft) which showed increased expression of hallmark stemness genes also showed increased expression of Notch1, LIF and PRKCA activity and increased chemoresistance to cisplatin, accompanied by decreased EZH2 activity and decreased H3K27me3 mark, further verifying that 1.4x A soft scaffolds were harboring the OS CSC population. The increased EZH2 and H3K27me3 levels in the populations from 1.4x NA stiff scaffolds which showed overall decreased stemness markers and increased cellular stiffness, ties in very well with a previous study (Zou et al., 2020) showing that increased H3K27me3 levels contributes to increased nuclei stiffness and led to H3K27me3-induced osteogenic differentiation in human bone marrow MSCs, further providing evidence linking H3K27 methylation with cell stiffness and stemness.

Doxorubicin resistance in CSCs is associated with the overexpression of drug efflux pumps, especially ABCB1, also known as P-glycoprotein (Lee et al., 2017). Consistent with the upregulated trend of other stemness properties and cisplatin chemoresistance in soft 1.4x A scaffold populations, we found that soft 1.4x A cell populations showed increased chemoresistance to Doxorubicin as compared to stiff 1.4x NA cell populations due to an increased expression of ABCB1 pump as determined through increased gene expression and enhanced protein expression observed through immunofluorescence of scaffold cultures. Interestingly, Notch1 has been reported to upregulate the expression of ABC transporters in CSCs (Cho et al., 2011), and we observed an increased expression of Notch1 and ABCB1 in doxorubicin and cisplatin-resistant soft 1.4x A subpopulation, indicating that mechanical cues might be linked to Notch/ABC-transporter mediated chemoresistance in mechanically induced CSCs. This agrees with reports from a previous study showing that Cisplatin resistance in OS CSCs was Notch-mediated (Yu et al., 2016). Moreover, the Notch pathway has been shown to be involved in regulating ALDH1A1 expression in OS (Mu et al., 2013), which is a strong contender of OS CSC marker (Mandell et al., 2022), that we also found to be increased in soft 1.4x A in conjunction with Notch1. However, the status of ALDH1A1 is debatable as contradictory evidence showed that their high expression was linked to better overall survival in OS patients (Haffner et al., n.d.). Similarly, other contenders of stemness markers such as CD44 and CD24 are the subject of ambiguity and debate because of inconsistency in expression across studies (Jaggupilli & Elkord, 2012).

Our RNA sequencing data showed 87 common genes with the same magnitude and direction of response to scaffold type in both cell lines. Some of the interesting findings were the enhancement of RAD50, and oncogenic long non-coding RNA SNHG1 (Small nucleolar RNA host gene 1) in soft 1.4xA cultures. RAD50 acts together with MRE11 and NBS1 in the MRN complex to fix DNA double-strand breaks and its enhanced activity promotes chemoresistance in tripe-negative breast cancer stem cells (Abad et al., 2021). Importantly, RAD50 is also involved in somatic stem cell reprogramming and increases DNA demethylation to establish induced pluripotent stem cells (Park et al., 2020). This suggests that exploring RAD50 in OS as marker of resistance and stemness might be of value in the future. SNHG1 has shown to be upregulated in OS has been linked to poor overall survival and increased tumor size by activating the WNT2B/Wnt/β-catenin pathway (Jiang et al., 2018) and by upregulating oncogene NOB1 (J. Wang et al., 2018). Most interestingly, SNHG1 has been reported to interact with members of the KLF family to regulate differentiation and tumorigenesis. Further investigation using qPCR determined that SNHG1 was indeed significantly increased in A soft culture as compared to NA stiff culture in both 143B and MG63.2. Members of the KLF family of transcription factors (Krüppel-like factors) play a complex role in many cancers (Z.-Y. Li et al., 2023) and many were differentially expressed between the two niches across both cell lines including KLF7/6/5/4/2, with KLF6 showing the highest expression difference (log fold change of 4, p-value 0.0001) across 143B and MG63.2 niche cultures. One study showed that SNHG1 helped maintain stem cell state and inhibited osteogenic differentiation by interacting with EZH2 to induce H3K27me3 methylation of KLF2 promoter, leading to KLF2 silencing (Z. Li et al., 2020). Another study showed that KLF4-induced upregulation of SNHG1 led to increased tumorigenesis in glioma through BIRC3 (Zhang et al., 2023). KLF7 has been shown to be overexpressed in OS (Huang et al., 2021) but the role of KLF6 is more complex, as its full and alternative splice variant (KLF6-SV1) can act either as a tumor suppressor or oncogene respectively (Hu et al., 2021). As the RNA sequencing data did not discriminate between the multiple variants of KLF6, we carried out separate qPCRs of KLF6 and KLF6-SV1 across both cell lines in the 2-scaffold models and determined that it was KLF6-SV1 that was significantly enhanced across 143B and MG63.2 in A soft niches as compared to NA stiff niches. Our study adds to the growing evidence of KLF6-SV1 as a target of interest implicated in OS stemness and warrants further investigation.

Not surprisingly, the RNA seq data also showed enrichment in enrichment in PPAR signaling pathway, ECM-receptor interactions and focal adhesion pathways, with the genes COL4A1 and ITGA3 showing higher expression in NA stiff groups across cell lines. Increased expression of COL4A1 has been linked to a more invasion tumor phenotype in carcinomas downstream of differentiation gene RUNX1-mediated expression, promoting invadopodia formation and triggering FAK-Src signaling (Miyake et al., 2017; Shin et al., 2022; T. Wang et al., 2020). . Integrin ITGA6, which has been implicated in OS stemness (Xu et al., 2025), was increased in the more stem-like population of 1.4x A soft scaffolds. However, ITGA3 has not been well studied in this context, but a specific polymorphism (rs2230392) in ITGA3 gene has been linked to increased OS incidence (Yang et al., 2014). A shortcoming of the current study is that it doesn’t explore the roles of integrin in detail, which would be crucial to study in future investigations to deeply understand the mechanotransductive pathways in stemness.

Our study showed that soft niche populations gave rise to greatest *in vivo* tumor growth and *in vitro* colony formation over stiff niche populations regardless of alignment. However, anisotropic (aligned) pores induced changes in tumor initiating properties within stiffer scaffolds, as evidenced by cells in A stiff scaffolds showing greater tumorigenicity than cells in NA stiff scaffolds in both *in vitro* colony assays and *in vivo* para-tibial tumors. A study by Janjanam et al. (Janjanam et al., 2021) demonstrated that microarchitectural anisotropy and Collagen I linearization in primary tumor sites led to increased metastases in breast cancer and it was mechanistically caused by WISP1 protein. Even though WISP1 has been linked to OS migration and angiogenesis and strongly correlates with tumor stage (Wu et al., 2013), its role in OS stemness is not clearly understood, and could be a valuable parameter to evaluate in future studies. Other studies using nanofabricated ECM, tissue elastography and X-ray computed tomography have also shown that tissue anisotropy and collagen fiber alignment provide directional and migration cues for metastatic growth in breast cancer and melanoma (Conceição et al., 2024; Kim et al., 2018; Leng et al., 2021).

Our study provided valuable insight into how macrophage to osteoclast activation might be driven by spatial heterogeneity across bone while also being implicated in tumor plasticity, with A soft showing higher active osteoclastogenesis along with increased stemness. Osteoclasts are the main drivers of bone resorption and remodeling, and their activation is highly significant in understanding primary OS and bone metastases, particularly in the context of bone region-specific tumor lesions and mechanotransductor protein interactions (Hirata et al., 2016). Future studies should build upon these findings and incorporate immune cells like T-cells considering that T-cell response is also biomechanically driven, with T-cell showing decreased cytotoxicity against soft cells, and cancer cells have been shown to use cellular stiffness as a mechanical immune checkpoint for T-cell evasion (Lei et al., 2021).

Our study underscores the importance of tumor heterogeneity and the need to carefully clarify what tumor subpopulations are being studied; oversimplification of behavior of the entire tumor might lead to missing nuances of how individual subpopulations might be behaving. Our findings suggest that niche heterogeneity in bone TME could give rise to cellular heterogeneity, which could be characterized by distinct gene profiles, plasticity and chemoresistant properties, suggesting that these subpopulations might be responding to therapies differently and working together to cause relapses and recurrence. Overall, we found a more stem-like population in soft, anisotropic bone niche as compared to the stiff, isotropic bone niche. The list of CSC biomarkers and pathways implicated in OS CSC development discussed in this study is not exhaustive, as biological systems are more complex than our current ability to model and understand them. Nevertheless, this study provides a new lens through which to look at the nuances of tumor heterogeneity and develop heterogeneity-driven therapeutic approaches, which could be broadly applied in the field of comparative oncology and towards bone metastases.

## METHODS

### Experimental Design

The schematic (Figure 8) summarizes the methodology of this study. 3D mineralized collagen scaffolds mimicking bone niches were used to mimic two mechanically and structurally distinct bone niches. These scaffold models were seeded with same human OS cell lines to evaluate the influence of mechanical forces on OS biology, with a focus on OS CSCs. Parameters for stemness evaluated were broadly divided into the categories of stemness genes, chemoresistance, and methylation. Further evaluation was done by conducting RNA-sequencing and testing cell stiffness of cell populations grown in these two distinct niches and doing *in vivo* tumorigenicity experiments of scaffold-harvested cells.

### Production of 3D mineralized collagen scaffolds

All mineralized collagen scaffolds used in this project were created at Professor Brendan Harley’s lab at University of Illinois Urbana-Champaign (UIUC). All 2-scaffold models were prepared by Aleczandria Tiffany, and all 4-scaffold models were prepared by Kyle Timmer as described briefly here. 3D porous mineralized collagen scaffolds with 47% collagen by weight were prepared by the freeze-drying method (Dewey et al., 2020). A slurry of bovine fibrillar collagen was prepared by blending dry collagen (8.1152g) in phosphoric acid (0.1456M) and calcium hydroxide (0.037M) buffer solution. Solutions of chondroitin sulfate (3.521g in 42.94 ml mineral buffer) and calcium hydroxide and calcium nitrate tetrahydrate (1.92g CaOH and 1.17g calcium nitrate tetrahydrate in 15 ml DI water) were prepared separately and mixed with hydrated collagen slurry.

The following equation was used to calculate collagen content based on weight percentage:

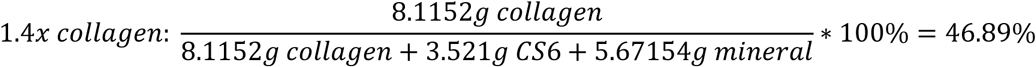

Mineralized collagen slurry was shaped into circular discs of 15 mm diameter and 3-4 mm height using molds and then processing them through a freeze-drying process as described previously (Dewey et al., 2020). Molds made of different materials were used to make scaffolds with different pore alignments. Scaffolds with non-aligned pores (NA scaffolds) and isotropic structure were made by using an aluminum mold, and scaffolds with aligned pores (A scaffolds) and anisotropic structure were made by using a Teflon mold with a copper base.

### Cell culture and seeding of scaffolds

Human osteosarcoma (OS) cell lines MG63.2, 143B, HOS and MNNG-HOS were grown in DMEM medium supplemented with 10% FBS and 1% penicillin/streptomycin at 37°C and 5% CO_2_.

To prepare scaffolds for cell culture, they were hydrated and crosslinked as described before (Dewey et al., 2020). Scaffolds were soaked in 70-100% ethanol, washed with PBS, crosslinked with EDC-NHS and washed with PBS again before being soaked in 1 ml OS-growth media for 48 hours in a 24-well plate. For 4-scaffold models, crosslinking ratios were varied to change stiffness of scaffolds, using the formula given below, where average weight of scaffolds was ∼29 mg. 1 ml of total crosslinking solution was used per scaffold.

<u>For soft scaffolds:</u>

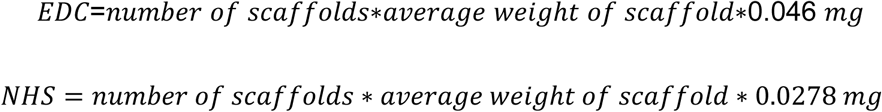

<u>For stiff scaffolds:</u>

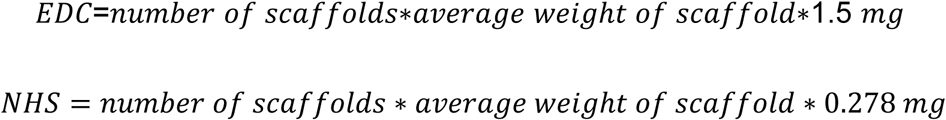

Soft A scaffolds from the 4-scaffold models are equivalent to 1.4x A scaffolds from the 2-scaffold models, and this group will be referred to as “soft A”, “soft 1.4x A” or “1.4x A” throughout the document. Similarly, stiff NA scaffolds from the 4-scaffold model are equivalent to 1.4x NA scaffolds from the 2-scaffold model, and this group will be referred to as “stiff NA”, “stiff 1.4x NA” or “1.4x NA” throughout the document.

Highly metastatic cell lines 143B and MG63.2 were used for seeding scaffolds in separate triplicate experiments. Before cell seeding, scaffolds were removed from the regular 24-well plate and transferred to an ultra-low attachment 24-well plate (Corning #CLS3473) without adding any media to the wells. Confluent culture of 143B and MG63.2 was collected using trypsin and centrifugation and counted. A total of 5×10^5^ to 2×10^6^ cells were seeded on scaffolds in separate replicate experiments; half of the cell suspension was pipetted on either side of the scaffold. Cells were allowed to attach on one side of the scaffold for 30 min before flipping the scaffold and seeding them on the other side. Cells were further allowed to attach for 1-2 hours before adding 1 ml OS-media to wells. 143B cells were allowed to grow on aligned and non-aligned scaffolds for 4 days before processing them for downstream experiments.

### Co-culture in scaffolds

A co-culture system was created to study osteoclast formation by co-culturing human OS 143B cells and RAW macrophage cell line. Initially, co-culture was done in petri plates to optimize culture conditions and visually verify osteoclast formation by TRAP5b staining. Scaffolds were seeded with 10K 143B OS cells and 5K RAW macrophage cells and on day 4 of co-culture, cell culture media was collected for evaluating TRACP5b levels using ELISA (Immunodiagnosticsystems #SB-TR103).

### Cell activity assessment in scaffolds

Cell viability and activity was assessed using alamarBlue and CellTiter-Glo. AlamarBlue was added 10% by volume to each well containing scaffold in media and incubated for 2 hours at 37°C. Each scaffold was completely submerged in total 1 ml volume of AlamarBlue-media in one well of a 24-well plate, covering both the top and bottom surfaces of the scaffold. Cell activity was assessed by measuring absorbance at 570 and 600 nm and calculating the percentage in reduction using the equation provided by Bio-Rad,

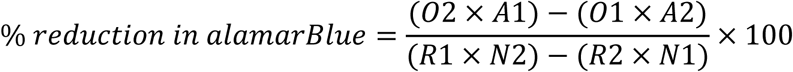

Cell activity in scaffolds using CellTiter-Glo (Promega #G7570) was assessed using 2-fold cell titrations in duplicate before using this assay for post-chemotherapy assays.

### Mechanical characterization of 3D scaffolds

#### Bulk modulus of scaffolds

Bulk compressive modulus of scaffolds was measured by doing compression tests with 5N load using Instron at the Carl R. Woese Institute for Genomic Biology (IGB) at UIUC. Compression test was performed on seeded scaffolds on day 4 of culture. Curve fitting was done using Matlab. Compression tests were performed with Aleczandria Tiffany and Kyle Timmer.

#### Atomic Force Microscopy (AFM)

##### AFM of dry scaffolds

AFM of dry scaffolds was carried out using the MultiMode AFM at the Beckman institute at UIUC. Gwyddion was used to process the AFM data.

##### AFM of hydrated, unseeded scaffolds

Force mapping of hydrated, unseeded scaffolds was carried at the Material Research Lab (MRL) at UIUC, with training provided by Kathy Walsh. Scaffold samples were fixed on a petri dish lid using 5-minute epoxy before placing in the AFM chamber and then completely submerged in PBS. The Asylum Research MFP-3D-SA was used for scanning and force mapping the scaffold surface in fluid. Two types of probes were used for force mapping; a sharp tip probe representative of integrin-scale mechanosensing, and a spherical probe representative of a whole cell-mechanosensing.

Initially, the ElectriCont-G probe with Chromium/Platinum coating from BudgetSensors was used in contact mode for all scans of hydrated A and NA scaffolds. The probe has a rectangular cantilever with a sharp pyramidal tip of 25 nm radius, with a force constant of 0.2 N/m and resonance frequency 13 kHz. The probe was first calibrated in air and then in fluid; a spring constant of 174.93 pN/m was obtained. A scan rate of 0.6 Hz and trigger point of 1-3 nN was used. Force maps of 10×10 um with increasing resolutions of 6, 10, 20, 256 pixels were taken. Corresponding height scans of same area were taken in contact mode. Force curves were fit to Hertz or JKR model as recommended by Asylum software.

The CP-qp-CONT-BSG-B-5 probe was purchased from NanoAndMore and was used in contact mode for force mapping all hydrated and unseeded A and NA scaffolds in fluid. The spherical tip of this probe had a diameter of 10 um and was made of borosilicate glass with a reflective chromium/gold plating. The beam cantilever of this probe had a force constant of 0.1 N/m and a resonance frequency of 30 kHz. The probe was first calibrated in air and then in fluid before performing force mapping in fluid. PBS buffer was the fluid of choice for submerging fixed scaffolds. A scan velocity of 3 um/s and trigger point of 5-15 nN was used. Force maps of 10×10 um were taken. Force curves were fit to the best fit model (JKR model) in the Asylum software. Averages of force curves were taken and compared to show differences in mechanics.

##### AFM of cells on scaffolds

Stiffness of cells growing on scaffolds was measured by contact mode force mapping in fluid using the CP-qp-CONT-BSG-B-5 probe and the Asylum MFP-3D. The scaffolds with live cultures were fixed in the center of the 10 mm petri-dish using superglue and allowed to attach for 15 minutes before adding media to the dish. The dish was placed on a place heater in the Bio-AFM-MFP-3D with the temperature set to 37C. After lowering the tip in the fluid, probe was calibrated and the virtual deflection, spring constant (100 pN/nm) and Invols were calculated. Force mapping was done using a trigger point of 1 nN, a scan rate of 0.17 Hz, scan velocity of 10 um/s, and XY velocity of 50 um/s. For the force map, a scan size of 50 um x 50 um was done across the scaffold surface by using a +/− 25 um offset in the X and Y axis. For the image scan of the same area, a scan rate of 0.5 Hz, set point of 0.3-0.5 V was used. Force maps of different resolutions were acquired, ranging from 32 to 96 scan points. JKR model in the Asylum software was used to fit the force curves. Parameters had to be manually adjusted to fit the JKR models and updated in the force map. Averages of force curves were taken and compared to show differences in mechanics.

### Environmental Scanning Electron Microscopy (ESEM) of scaffolds

Cultured scaffolds on day 4 of culture were fixed in 10% formalin on the shaker and then transferred to 70% ethanol. Scaffolds were sputter-coated and ESEM at the Beckman institute of UIUC was used for imaging the surface and cross-sections of unseeded and seeded scaffolds. Compression tests were performed with Aleczandria Tiffany.

### RNA extraction from scaffolds

Seeded scaffolds were removed from media-containing wells on day 4 of the culture and immersed in 2 ml TRIzol reagent (Invitrogen) per scaffold in a 5 ml tube and homogenized on ice using a tissue homogenizer. After complete pulverization of scaffold, the TRIzol-lysate was spun down at the highest speed to separate collagen debris from cell lysate. TRIzol-cell lysate (supernatant) was collected and filtered once through the Direct-zol RNA Miniprep column from Zymo research to further remove fine collagen debris – this step is crucial to obtaining high purity and high-quality RNA. The filtered lysate was then processed according to the Direct-zol RNA Miniprep protocol and RNA was eluted in 25 ul DNase/RNase free water. The quality and quantity of RNA was measured using the NanoDrop machine (Thermo Scientific #13-400-518). Only samples with the A260/280 ratio of 1.8-2.0 were used for downstream applications.

Scaffold-harvested cells were also grown for 10 days on standard plastic petri-dishes and then used for RNA extraction and qPCR of target genes to check if gene expression differences are maintained. RNA extraction from regular petri-dish 2D cultures were done by collecting cell pellets and processing with TRIzol reagent and Direct-zol RNA Miniprep kit.

### qPCR of target genes

Extracted RNA was reverse transcribed to cDNA using SuperScript IV Reverse Transcriptase and Oligo d(T)_20_ primer (Invitrogen #18091050). A final concentration of 2.5 ng/ul cDNA was obtained and used for real time PCR. Primers were created using Primer-BLAST (NIH) and ordered from IDT and Sigma-Aldrich. The primer sequences used for each target gene are given in Supplemental Table 1.

PowerUp SYBR green mastermix (Invitrogen #A25742) was used for all qPCR reactions. A total reaction volume of 20 ul per well in a 96-well reaction plate was made for each target gene. GAPDH was used as the housekeeping gene. Real time PCR settings provided by SYBR green protocol were followed to run all qPCR reactions. The 2^−ΔΔCt^ method was used to process qPCR data and statistical analysis was done on GraphPad using unpaired t-test after confirming normality of data. A p-value of less than 0.05 was considered significant.

### RNA-Sequencing of niche populations

Human osteosarcoma lines 143B and MG63.2 were seeded separately at a density of 5×10^5^ cells on 1.4x A and 1.4x NA scaffolds each and grown for 4 days. RNA was extracted on day 4 as explained above and was assessed using Nanodrop before being sent to the HPCBio core facility at the Roy J. Carver Biotechnology Center at UIUC. The core facility carried out the quality control assessment and sequencing of RNA samples extracted from scaffold cultures. Salmon version 1.10.0 was used by the core facility to quantify transcript expression from RNA-Seq data. Normalization was done using RUV analysis and sample clustering was performed. Pairwise comparison (A vs NA) was done for each cell line (143B, MG63.2) and interaction between cell lines was also analyzed.

### Protein expression and activity in scaffold populations

ELISA kits were used to assess activity levels of EZH2 (BPS Bioscience catalog# 52009L), PKC Kinase (abcam #139437) and LIF (Thermofisher #BMS242). Protein lysate extracted from cultured scaffolds was used for EZH2 and PRCKA assays because these are intracellular proteins, while scaffold media on day 4 was collected and used for LIF assay to assess levels of LIF secreted in the media.

### Immunofluorescence of 2D cultures and seeded scaffolds

143B, HOS, MNNG-HOS cells were grown on glass-bottom dishes, fixed in 4% paraformaldehyde for 10 min, blocked with NGS and stained for EZH2 (Cell Signaling #5246) and H3K27me3 (Cell Signaling #9733). Images were captured using confocal microscopy.

Cultured scaffolds on day 4 of culture were fixed in 10% formalin on the shaker and then transferred to 70% ethanol. Fixed scaffolds were embedded in paraffin, sectioned, and used for IHC-P and immunofluorescence. All embedding, sectioning and staining steps were carried out by the histology lab at the College of Veterinary Medicine. Cell attachment and growth on both sides of the scaffolds was confirmed by H&E staining. Anti-actin antibody and DAPI were used to visualize cytoskeletal changes by immunofluorescence. Protein expressions of P-glycoprotein (abcam #ab235954), ALDH1A1 (ThermoFisher #60171-1-IG), active YAP (abcam #ab205270), and H3K27me3 (Cell Signaling #9733) were visualized by immunofluorescence of FFPE scaffolds. ImageJ was used to quantify the corrected total cell fluorescence (CTCF).

### Chemosensitivity assays

Day 4 cultures of 1.4x A and 1.4x NA seeded scaffolds were treated with two-fold serial dilutions of doxorubicin (0.15 – 10 uM) for 72 hrs and cell viability was assessed using alamarBlue. Day 4 cultures of 1.4x A and 1.4x NA seeded scaffolds were treated with two-fold serial dilutions of cisplatin (1.5 – 200 uM) for 72 hrs and cell viability was assessed using alamarBlue. In another set of replicate experiments, day 4 cultures of 1.4x A and 1.4x NA seeded scaffolds were treated with two-fold serial dilutions of cisplatin (0.0625 – 16 uM) for 48 hrs and cell viability was assessed using CellTiter-Glo.

### Colony Assays

#### Soft agar colony assay

The soft agar method for colony forming assay is more representative of CSCs growing in a soft microenvironment similar to soft organs of metastatic tropism such as the lung. The assay was conducted in a 6-well plate using complete DMEM media (10% FBS, 1% Pen-Strep antibiotics) and low-melting temperature molecular grade Agar. 0.6% of agar media was prepared in a sterile glass bottle and microwaved to properly dissolve agar in complete media. A base layer of 1.5 ml of 0.6% agar media was first poured in each well and allowed to cool and solidify for 30 minutes at room temperature in a cell culture hood. Cells harvested from scaffolds were counted using an automatic cytometer and 1000 cells were resuspended in complete media. Cell suspension was mixed in a 1:1 ratio with the 0.6% agar media to make a final concentration of 0.3% agar media and 1.5 ml was poured on top of the base layer in each well. Top layer was allowed to solidify like the base layer and 100 ul complete media was added on top. Images of unfixed and unstained colonies were taken after 23 days using brightfield/phase contrast setting on a Keyence BZ-X800 microscope. Samples were plated in triplicates.

#### Standard colony assay

For 143B cells harvested from scaffolds, two colony assays were done at different cell densities in triplicates. 100 cells and 1000 cells were plated per well from each scaffold (1.4x A, 1.4x NA) in 6-well plates and grown for 13 days in complete media before fixing and staining with 0.6% glutaraldehyde and 0.5% crystal violet. For MG63.2 cells harvested from scaffolds, 1000 cells from each scaffold type were plated in a 6-well plate and grown for 15 days before staining with 0.6% glutaraldehyde and 0.5% crystal violet. All groups were plated in replicates of 6 in a single experiment and the experiment was replicated thrice at least. Colony assays were imaged using the iBright imager (Invitrogen) and analyzed using the iBright software.

### Mouse models

Human OS cells grown on collagen scaffolds were harvested on day 4 and used for creating OS xenograft tumors in nude athymic mice. Nude athymic mice were purchased from Jackson Laboratory. Scaffold cultures were washed with 1x PBS and then incubated in Liberase TL (Sigma #05401020001) diluted in serum-free media at a final concentration of 0.5 WU/ml for 1 hour on an orbital shaker at 37C and 5% CO_2_. Liberase-cell suspension was collected, and scaffolds were incubated in TrypLE for ∼20 minutes, followed by consolidation of cell suspensions released from Liberase and TrypLE treatments. Cell suspensions were neutralized with complete media and filtered multiple times through 40 um and 20 um cell strainers to remove as much scaffold debris as possible. Cells were counted in an automatic cell counter and washed and resuspended in 1x HBSS and mixed at a 1:1 ratio with Matrigel for para-tibial injection. All para-tibial injections were carried out by Dr Tim Fan. A final concentration of 1×10^6^ cells per 50 ul was injected into the para-tibial space along the right tibia. MG63.2 and 143B-scaffold culture tumors in mice were left to grow for 100 days and 60 days respectively before sacrifice and tissue collection, unless tumor diameter of 20 mm was reached, in which case the mice were sacrificed immediately according to IACUC protocol. Tibial tumors were measured and whole tibia with tumor along with corresponding lungs were processed as FFPE tissues for H&E staining. FFPE tissue processing and slide staining were done at the Histology lab at Vet Med or the TEP core at the Cancer Center at Illinois.

### Statistical Analysis

Statistical analysis was done on GraphPad using unpaired t-test or ANOVA or Kruskal-Wallis depending on the data set after confirming normality of data. A p-value of less than 0.05 was considered significant. All experiments were done in triplicates and minimum sample size of n=3 was used in all experiments. All PCR data was normalized to the housekeeping gene GAPDH.

## Supporting information

Supplemental Images and Table

