## Supplemental Images and Table for "TUMOR HETEROGENEITY AND STEMNESS CAN BE SHAPED BY MECHANO-STRUCTURAL PARAMETERS IN OSTEOSARCOMA"

### SUPPLEMENTAL MATERIAL

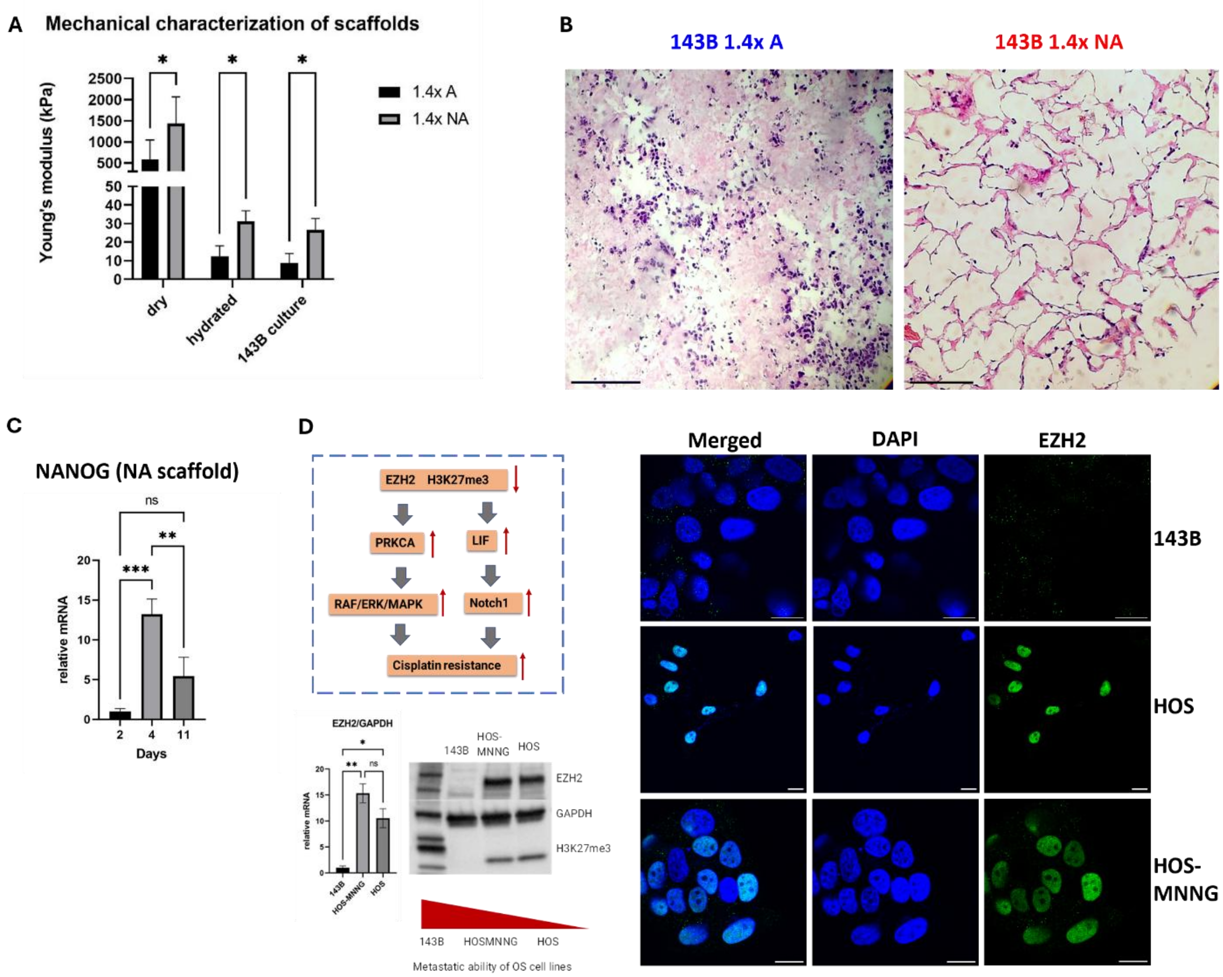


**Supplementary Figure 1. (A)** 2-scaffold models (1.4x A and 1.4x NA) have significantly different mechanical properties across all conditions (dry, hydrated or seeded), making them a robust model. **(B)** H&E of FFPE 143B cultures on soft 1.4x A and stiff 1.4x NA scaffolds, showing difference in cell attachment and penetration pattern across the two niches (scale bar=100um). **(C)** Time point analysis of gene expression in 143B NA scaffolds, showing that day 4 has peak NANOG expression, which is one of the hallmark stemness genes assessed in the study. **(D)** Schematic showing the pathways connecting methyltransferase EZH2 and H3K27me3 methylation with Cisplatin resistance in OS, as reported by He et al., 2019. Western blot, qPCR and super resolution immunofluorescence data (scale bar=20um) for the study was done to verify EZH2 and H3K27me3 expression in metastatic line 143B and non-metastatic line HOS.

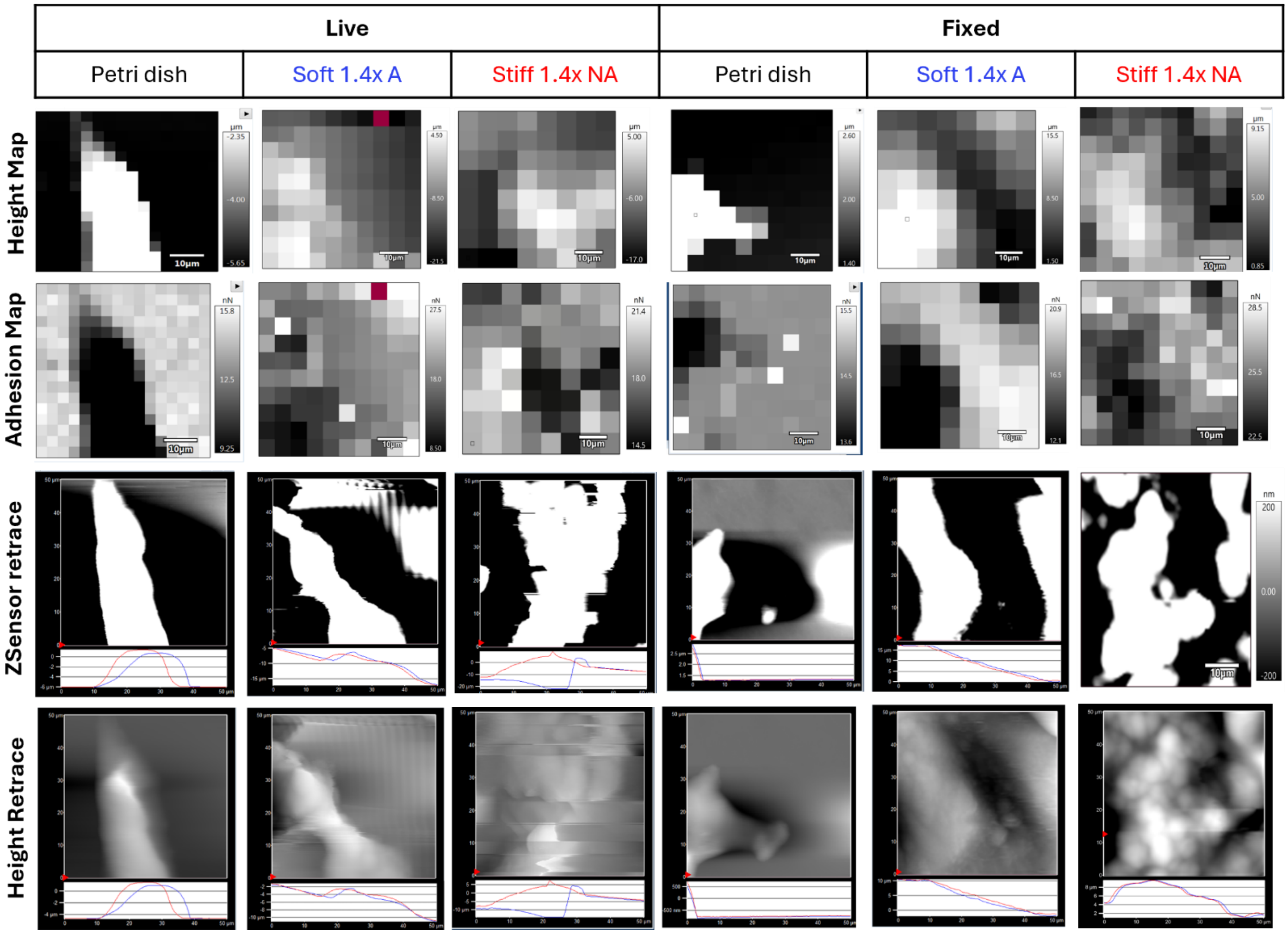


**Supplementary Figure 2. AFM data showing height map, adhesion map. ZSensor retrace and height retrace of live and fixed 143B cells on petri dish, soft 1.4x A and stiff 1.4x NA scaffolds.** These parameters show cell shape and help mark out the cell on petri dish and scaffold surfaces, as increased height corresponds to the cell. Although the scaffold surface is softer and more heterogeneous than plastic surface, the cell outline is not as sharp. Additionally, adhesion map further confirms that the force map scanned is of the cell, with adhesion and height of cell generally showing up in an inverse relationship on the graphs.

| Gene | Primer sequence (5' → 3') |
|---|---|
| GAPDH forward<br>GAPDH reverse | ACAGTCAGCCGCATCTTCTT<br>ACGACCAAATCCGTTGACTC |
| NANOG forward<br>NANOG reverse | CAATGGTGTGACGCAGAAGG<br>GCATCCCTGGTGGTAGGAAG |
| SOX2 forward<br>SOX2 reverse | AGCATGGAGAAAACCCGGTA<br>AGTGTGGATGGGATTGGTGT |
| OCT4 forward<br>OCT4 reverse | CCTGAAGCAGAAGAGGATCACC<br>TTGGCTGAATACCTTCCCAAA |
| ALDH1A1 forward<br>ALDH1A1 reverse | CACCAGGGCCAGTGTTGTAT<br>CTTAGCCCGCTCAACACTCC |
| EZH2 forward<br>EZH2 reverse | GACTCAGAAGGCAGTGGAGCC<br>AGTCTGGCCCATGATTATTCTTCGT |
| LIF forward<br>LIF reverse | CTTGGCGGCAGGAGTTGT<br>GTTGACAGGGGTGATGGGG |
| PRCKA forward<br>PRCKA reverse | TGCACGAGGTGAAGGACCA<br>TTTCCCAAACCCCCAGATGAA |
| KDM6A forward<br>KDM6A reverse | CTACCAATTCCCGCAGAGCTTAC<br>GGATGATGAGGTTTACATGCCTGC |
| KDM6B forward<br>KDM6B reverse | CCTTCACGCTGGCCCC<br>AGGCATGTGCTGGTGGC |
| ABCB1 forward<br>ABCB1 reverse | CGGTTTGGAGCCTACTTGGT<br>ATGAACTGACTTGCCCCACG |
| FAK1 forward<br>FAK1 reverse | CAACGAGGGTGTCAAGCCAT<br>GTCCAGGTTGGCAGTAGGAG |
| YAP1 forward<br>YAP1 reverse | GAACTGCTTCGGCAGGCAAT<br>GTGTTGGTAACTGGCTACGC |
| TAZ forward<br>TAZ reverse | CAATCTCGGGACTCACCCG<br>CGGATTCATCTTCTGGGCGG |
| Notch1 forward<br>Notch1 reverse | ATGCCTGCCTCACCAACC<br>GGCACGATTTCCCTGACCA |
| RANKL forward<br>RANKL reverse | CTTTCAAGGAGCTGTGCAAAAGG<br>GCCATCCACCATCGCTTTCTC |
| PRCKA forward<br>PRCKA reverse | TGCACGAGGTGAAGGACCA<br>TTTCCCAAACCCCCAGATGAA |
| SNHG1 forward<br>SNHG1 reverse | TGGTAAGTGGCTTCGTGGTC<br>GCGTGCTATGCTATGTTCGC |
| KLF6 forward<br>KLF6 reverse | GTGCCACTTTAACGGCTGC<br>TCTGTAAGGCTTTTCTCCTGTGTG |

**Supplemental Table 1. Sequences for qPCR primers used in the study.**

| Gene | Primer Sequence (5' → 3') |
|---|---|
| KLF6-SV1 forward<br>KLF6-SV1 reverse | CCTCCACGCCTCCATCTTCT<br>TAAGGCTTTTCTCCTTCCCTGG |
| KLF4 forward<br>KLF4 reverse | CGAACCCACACAGGTGAGAAA<br>ACGGTAGTGCCTGGTCAGTT |
| ITGA6 forward<br>ITGA6 reverse | AATGCAGGCACTCAGGTTCG<br>ATCCACCAAGGTACTCCCGA |
| ITGB1 forward<br>ITGB1 reverse | ATCCACCAAGGTACTCCCGA<br>ATCCACCAAGGTACTCCCGA |
| TRACP5b forward<br>TRACP5b reverse | GACCCACCGCCAAGATGGAT<br>GGTCAGGAGTGGGAGCCATA |

**Supplemental Table 1 (continued). Sequences for qPCR primers used in the study.** All sequences are human specific except for TRACP5b which is murine specific.
